# Genome-Wide Mapping of Major Histone Modifications Reveals Distinct Epigenetic Regulatory States in a Reef-Building Coral

**DOI:** 10.64898/2026.09.10.745915

**Authors:** George Warfel, Shanna L. Davidson, Luke Sadalla, Tooba Varasteh, Harun Ozturk, Fidan Seker-Polat, Vadim Backman, Mazhar Adli, Luisa A. Marcelino

## Abstract

Reef-building corals exhibit extensive physiological and transcriptional plasticity, yet the chromatin-level regulation of coral gene expression remains poorly characterized. Here, we generated the first genome-wide maps of major histone modifications in a reef-building coral by performing chromatin immunoprecipitation sequencing (ChIP-seq) in adult *Pocillopora damicornis* maintained under ambient conditions. We profiled two active marks, H3K4me3 and H3K27ac, and two repressive marks, H3K27me3 and H3K9me3, and integrated these maps with RNA sequencing (RNA-seq) to relate chromatin state to transcriptional output. As in other eukaryotes, H3K4me3 and H3K27ac were enriched around transcriptional start sites and positively associated with gene expression, whereas H3K27me3 and H3K9me3 showed broader enrichment patterns and were negatively associated with transcription. Partitioning highly expressed genes by promoter chromatin state revealed that promoters with strong H3K4me3/H3K27ac signal carried YY1-family motifs and served core cellular functions, while a second group enriched for cell-surface receptors and proteolysis lacked these active promoter marks and motif repertoire, despite similar transcript abundance. Repressive marks also distinguished functional genomic states: H3K27me3 was enriched over lowly expressed receptor-like and developmental gene classes, whereas H3K9me3 was prominent at transposable element-like loci and broad non-genic regions. We further identified H3K27ac-enriched, H3K4me3-low candidate enhancer-like elements proximal to core promoters, many near genes with transcription factor-related functions. Together, these results define a baseline chromatin-state landscape in reef-building corals, offering a resource for validating targeted chromatin assays and a foundation for future studies of coral gene regulation across environmental, developmental, and symbiotic contexts.

**Graphical Abstract:** 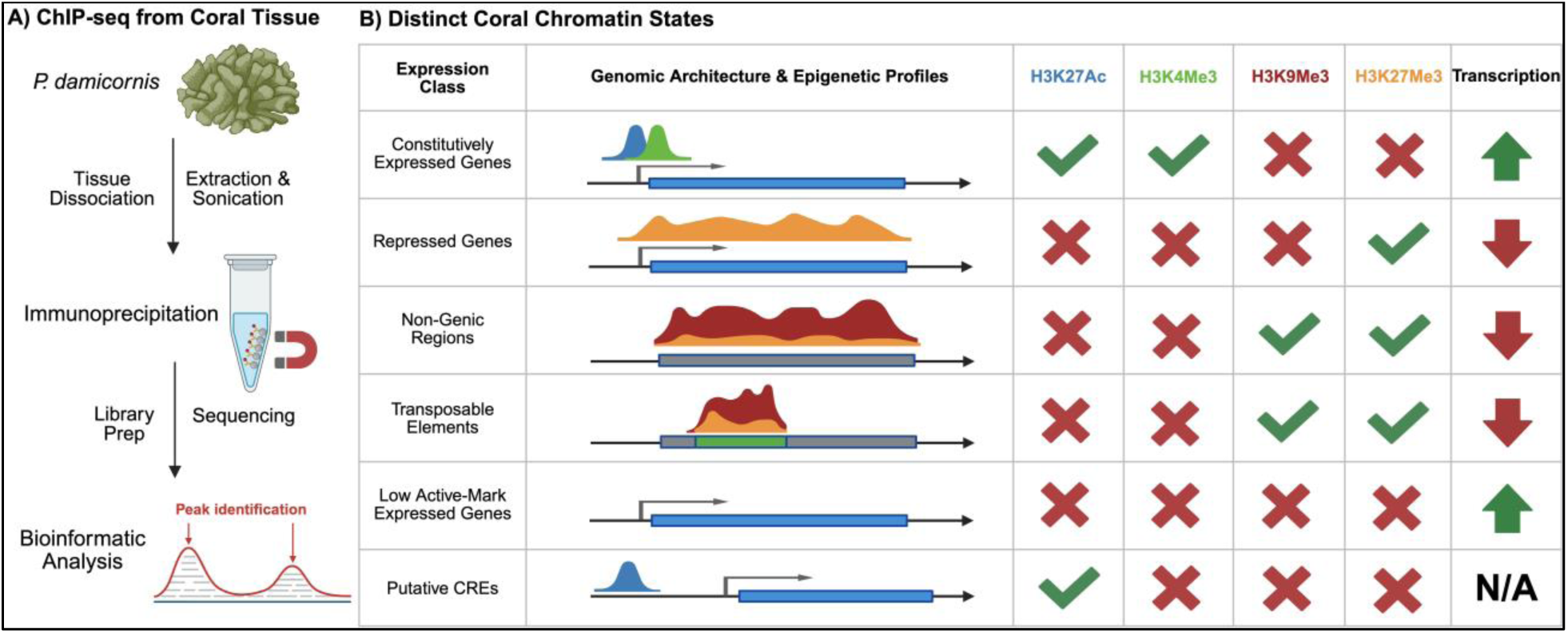

## Introduction

Coral reefs support marine biodiversity, provide food for millions of people, and protect coastlines, but increasingly frequent and severe marine heatwaves have driven widespread losses in coral abundance, diversity, and reef function, with some dominant reef-building corals now reaching functional extinction in heavily impacted regions (Eddy et al., 2021; Hughes et al., 2018; Manzello et al., 2025; Skirving et al., 2019). Coral responses to heat stress are nonetheless highly variable among species, populations, and individuals (Humanes et al., 2022; Kenkel & Matz, 2016; Voolstra et al., 2021). This has been attributed to multiple interacting mechanisms, including host genetic variation, algal symbiont composition, microbiome dynamics, environmental history, physiological acclimatization, transcriptomic plasticity, and epigenetic regulation (as reviewed in Voolstra et al., 2021).

Much of this variation has been measured as changes in gene expression, including transcriptional plasticity, frontloading, dampening of transcriptional responses, and transcriptomic resilience, and has been linked to acclimatization and thermal tolerance (Barshis et al., 2013; Drury et al., 2022; Palumbi et al., 2014; Savary et al., 2021; Stick et al., 2026). However, transcript abundance is the outcome of upstream regulatory processes that determine which genes are accessible, active, or repressed in a given context. Interpreting how these gene expression programs shift under stress requires knowing how they are organized at baseline, such as which genomic features carry regulatory information and how those features relate to gene activity, but this architecture remains largely uncharacterized in corals.

Epigenetic mechanisms provide a route by which environmental variation can alter gene-expression states without changing the DNA sequence. Across marine organisms, DNA methylation, histone modifications, chromatin accessibility, histone variants, and noncoding RNAs have been implicated in phenotypic plasticity, acclimatization, development, stress responses, and biomonitoring, although their interactions remain poorly resolved (Brunner et al., 2026; Eirin-Lopez & Putnam, 2019). In corals, most epigenetic work has focused on DNA methylation, linking methylome variation with stress responses (Guerrero & Bay, 2024; Liew et al., 2018; Putnam et al., 2016), seasonal and spatial environmental variation (Hackerott et al., 2023), and light-induced phenotypic plasticity (Gomez-Campo et al., 2024). Histone modifications, by contrast, remain understudied in corals.

Among epigenetic mechanisms, histone post-translational modifications (PTMs) are central regulators of chromatin state because they influence DNA accessibility, recruit regulatory complexes, and mark genomic regions associated with transcriptional activation or repression (as reviewed in Millán-Zambrano et al., 2022). Acetylation of histone tails, which neutralizes lysine residues and weakens DNA-histone interactions, is generally associated with higher transcriptional activity, while histone methylation has more complex regulatory consequences depending on the residue and degree of methylation (Bannister & Kouzarides, 2011; Smolle & Workman, 2013). In many eukaryotes, H3K4me3 and H3K27ac are associated with active promoters and regulatory regions, whereas H3K27me3 and H3K9me3 mark distinct forms of repressive chromatin (Igolkina et al., 2019; Kim & Kim, 2012).

Genome-wide chromatin studies in early-branching animals suggest that these regulatory roles are deeply conserved. In the sponge *Amphimedon*, histone modification maps revealed conserved features of animal cis-regulatory complexity (Gaiti et al., 2017). In the coral model *Aiptasia*, histone modifications, DNA methylation, and chromatin accessibility act together to regulate symbiosis and stress-related gene expression programs (Li et al., 2018; Nawaz et al., 2022; Weizman & Levy, 2019). In *Nematostella*, chromatin profiling has identified conserved promoter and enhancer-like regulatory landscapes associated with developmental gene regulation (Schwaiger et al., 2014). Yet, how far these features extend to reef-building corals remains unknown. Does the presence of active histone modifications consistently predict transcript abundance? Do canonical repressive modifications mark distinct genomic contexts as they do in bilaterians? And do coral cis-regulatory elements carry sequence-encoded regulatory information, in a genome markedly more compact than those of complex bilaterians?

Addressing these questions requires genome-wide, position-resolved measurements that are not yet available for corals. Recent transcriptomic work shows that expression of histone variants and chromatin modifier enzymes changes markedly during coral development (Brunner et al., 2026), and mass-spectrometry profiling has begun cataloguing coral histone modifications and their abundance under heat stress (Fuller et al., 2025a; Fuller et al., 2025b). Together these establish that coral chromatin machinery is dynamic and that histone modifications are present and quantifiable, but neither approach can resolve where in the genome these states are deposited.

Here, we optimized chromatin immunoprecipitation sequencing (ChIP-seq) for adult *Pocillopora damicornis* grown under ambient conditions and generated genome-wide maps of four histone modifications associated with active and repressive chromatin states: H3K4me3, H3K27ac, H3K27me3, and H3K9me3. We integrated ChIP-seq with RNA sequencing (RNA-seq) to test how histone modifications are associated with gene expression and whether active and repressive marks identify distinct gene classes or chromatin states. By profiling both active and repressive histone modifications, this study establishes a baseline chromatin landscape in reef-building corals and provides a foundation for future studies of chromatin remodeling across environmental, developmental, and symbiotic contexts.

## Results

### Genome-wide chromatin state maps of major histone modifications reveal key regulatory elements in *Pocillopora damicornis*

To compare genome-wide distributions of histone modifications, we computed ChIP-seq signal for each library in 1kb genomic bins. After hierarchical clustering and Spearman correlation analysis, three distinct clusters emerged: Activating marks H3K4me3 and H3K27ac, repressive marks H3K27me3 and H3K9me3, and input controls (Figure 1A). Replicate libraries for each histone modification exhibited strong correlation (ρ=0.82-0.94), whereas input controls were moderately correlated (ρ=0.66). Together with antibody validation (Figure S1), these results support the reproducibility and biological interpretability of the ChIP-seq datasets.

**Figure 1.**
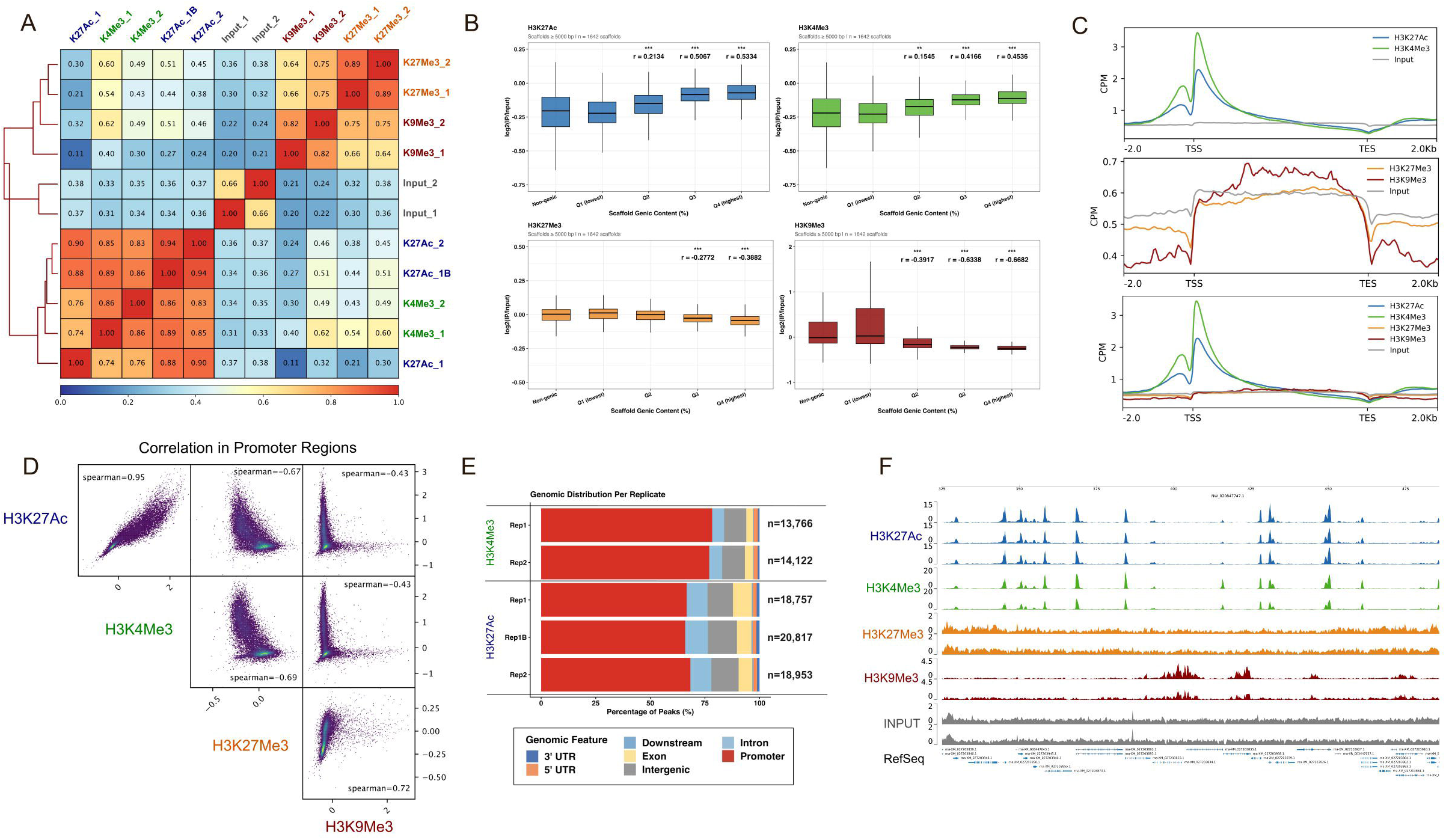
Genome-wide chromatin state maps of major histone modifications reveal key regulatory elements in *Pocil/opora damicornis*. A) Spearman correlation heatmap with hierarchical clustering of ChIP-seq libraries using 1kb genomic bins, after greylist filtering (red: high correlation, blue: low correlation). B) Mean log2(1P/input) enrichment across scaffolds ranked by gene density. All scaffolds > 5 kb were classified as non-genie or divided into four gene-density quartiles (Q1, low; Q4, high gene density). Significance and effect sizes were determined using SH-corrected Wilcoxon rank-sum tests comparing non-genie scaffolds with each genie scaffold group. C) CPM-normalized ChlP-seq coverage profiles across RefSeq-annotated genes. D) Pairwise correlations of mean log2(1P/input) signal across RefSeq promoters (±1kb from TSS), after greylist filtering. E) Distribution of macs2-identified H3K4me3 and H3K27ac peaks across genomic features for all replicates (isogenic replicates Rep1 and Rep2; technical replicates Rep1 and Rep1B), annotated with *Ch/Pseeker.* F) ChIP-seq tracks across a representative >150kb genomic region; replicates are shown as separate tracks.

We next examined whether histone modification enrichment varied with scaffold gene density. Activating marks H3K4me3 and H3K27ac exhibited higher mean ChIP signal on gene-dense scaffolds than on non-genic scaffolds (Figure 1B, BH-corrected Wilcoxon rank-sum). In contrast, repressive H3K9me3 signal was highest on non-genic and low gene-density scaffolds and decreased across higher density scaffolds. H3K27me3 also demonstrated higher ChIP signal in non-genic scaffolds versus the top 50% most gene-dense scaffolds, but with much lower effect size (r= −0.277, −0.388 vs. r= −0.634, −0.668 for H3K9me3).

We then mapped histone modifications within annotated NCBI *Pocillopora damicornis* RefSeq genes. H3K4me3 and H3K27ac were sharply enriched around transcription start sites (TSS), while H3K27me3 and H3K9me3 were depleted in the regions flanking genes, and showed broader, lower-amplitude signal across gene bodies (Figure 1C). These divergent profiles highlight distinct gene-associated distributions of active and repressive histone modifications. Note that input coverage also increased over gene bodies, likely reflecting preferential fragmentation of open and actively transcribed chromatin during sonication (Auerbach et al., 2009). Thus, Figure 1C shows CPM-normalized coverage with the input track displayed for reference, but all quantitative analyses use log2(IP/input) normalization to control for this artifact.

We next compared mean ChIP signal across annotated promoters (TSS ±1 kb) and tested pairwise correlations between marks. H3K4me3 and H3K27ac were strongly correlated in promoters (ρ=0.95), as were H3K27me3 and H3K9me3 (ρ=0.72; Figure 1D), indicating that the two repressive marks co-occur at promoters despite differing in their scaffold-level distributions (Figure 1B). In contrast, activating marks (H3K4me3 and H3K27ac) were negatively correlated with H3K27me3 (ρ= −0.69 and −0.67), and more weakly with H3K9me3 (ρ= −0.43 and −0.43). Genomic annotation of identified peaks showed that most H3K4me3 (n=13,766 and 14,122 across replicates) and H3K27ac (n=18,757, 20,817 and 18,953) peaks occurred near promoters, but H3K4me3 had slightly higher promoter dominance than H3K27ac (Figure 1E).

Genome browser tracks further illustrate these trends: H3K4me3 and H3K27ac occurred in sharp peaks in promoters, while repressive modifications had broader peak patterns (Figure 1F). Genes with strong active-mark promoter peaks generally showed lower repressive mark signal across gene bodies. In contrast, gene bodies enriched in repressive marks showed weak promoter signal for active marks.

### Epigenomic states of regulatory elements are linked to gene expression programs

To examine links between histone modifications and gene expression, we visualized ChIP signal across RefSeq genes, which were classified as undetected by RNA-seq or divided into four expression quartiles. These analyses revealed that H3K4me3 and H3K27ac enrichment around the TSS increased across expression quartiles, from the lowest- to highest-expressed genes (Figure 2A; Figure S2; Table S1; p<0.001 Kruskal Wallis & Dunn’s test). While highly expressed genes had strong active mark peaks, undetected genes or those in the lowest expression quartile (Q_1_) had lower signal of H3K4me3 and H3K27ac. In contrast, H3K27me3 and H3K9me3 were highest in undetected genes and Q1 genes relative to Q2-Q4 genes (p<0.001, Kruskal Wallis & Dunn’s test; Figure 2A; Table S1), with H3K9me3 highest in undetected genes (Figure S2; Table S1; p<0.001, Kruskal Wallis & Dunn’s test). Both repressive marks decreased with increasing gene expression, especially around the TSS. Overall, activating and repressive modifications displayed opposing patterns relative to gene expression.

**Figure 2.**
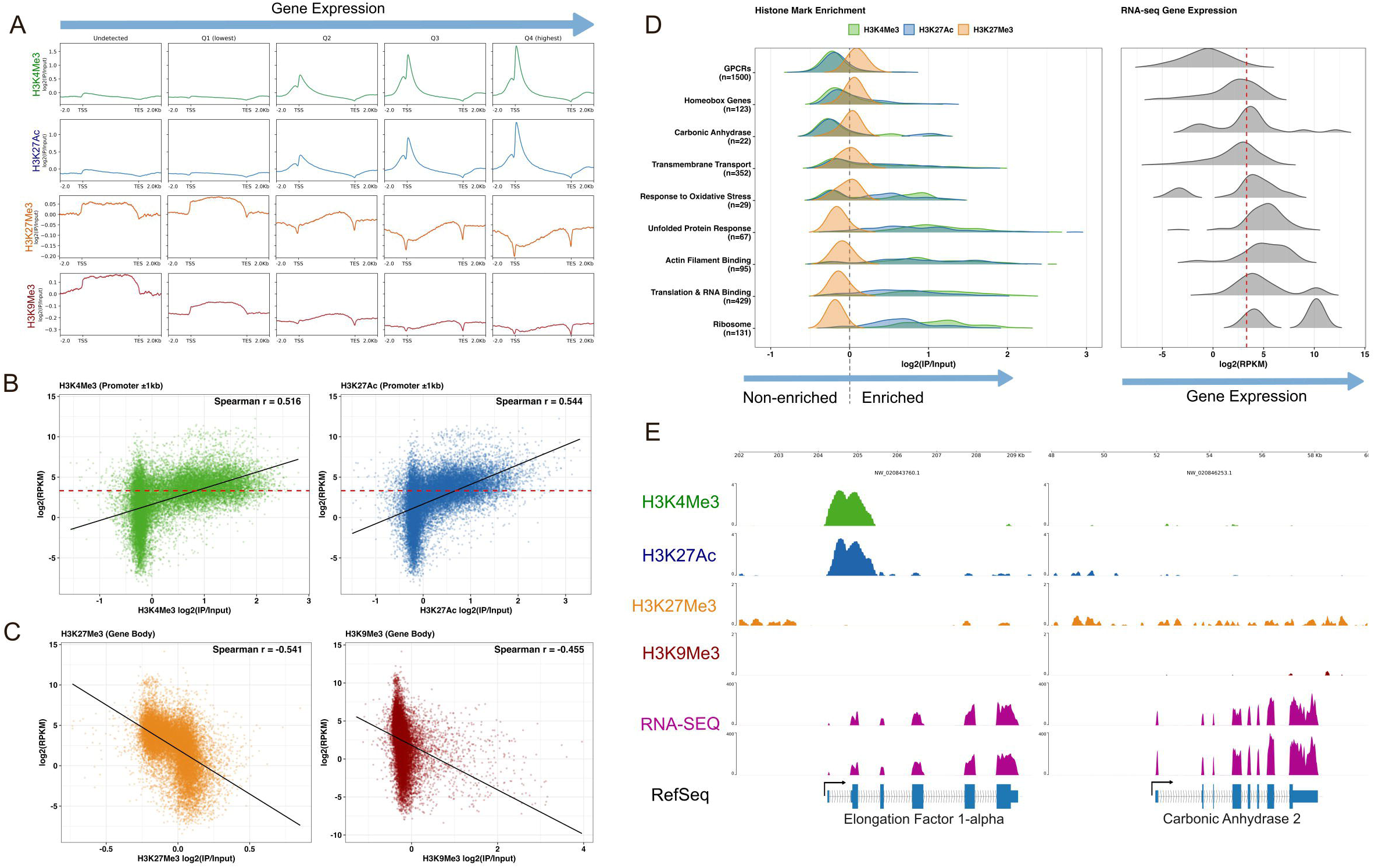
Epigenomic states of regulatory elements are linked to gene expression programs. A) Log2(IP/input) profiles across RefSeq genes grouped as undetected (n=1,546) by RNA-seq or divided into four expression quartiles among detected genes, with increasing RPKM values from left to right (n= 5,297 per quartile). B,C) Relationship between log2(RPKM) expression and log2(IP/input) ChlP signal for genes detected by RNA-seq (n=21,189). B) H3K4me3 and H3K27ac scores were computed across promoters (±1kb from TSS). C) H3K27me3 and H3K9me3 scores were computed across gene bodies (TSS-TES). Red dashed lines denote RPKM > 10. D) Distributions of active (K4me3 + K27ac) promoter signal, repressive K27me3 gene-body signal, and RNA-seq gene expression for selected gene groups. Black dashed line indicates log2(IP/input) = 0; red dashed line indicates RPKM > 10. E) log2(IP/input) ChIP-seq and RNA-seq tracks across Elongation Factor 1-alpha (RPKM=2,654) and Carbonic Anhydrase-2 (RPKM=4,404), two highly expressed genes with divergent chromatin profiles.

Next, we compared histone modification enrichment with RNA-seq expression for all detected RefSeq genes (n=21,189, RPKM > 0). Promoter H3K4me3 and H3K27ac scores (TSS ±1kb) were positively correlated with gene expression (Spearman ρ= +0.516, +0.544 respectively; Figure 2B). However, a subset of highly expressed genes (RPKM > 10) lacked strong promoter enrichment of active marks, which we examine below. Because the repressive marks were distributed across gene bodies rather than concentrated at promoters, we computed their scores over gene body (TSS-TES) intervals. Both were negatively correlated with expression (Spearman ρ = −0.541, −0.455 respectively; Figure 2C), yet their score distributions differed (H3K27me3 signal spanned roughly −0.6 to +0.8, while H3K9me3 spanned −1 to +4, with a long positive tail), suggesting distinct repressive chromatin states for these two histone modifications.

We then asked whether these chromatin patterns differed amongst functional gene classes. Gene groups with higher H3K27me3 and lower H3K4me3/H3K27ac signal generally showed lower expression (Figure 2D). G-protein coupled receptors (GPCRs) exhibited the clearest repressive chromatin state and the lowest median expression among the groups examined. Homeobox, carbonic anhydrase-related, transmembrane transport, and oxidative stress-related genes showed broader or multimodal chromatin and expression distributions. In contrast, unfolded protein response genes, and constitutive genes involved in actin binding, translation, and ribosome function showed highly active chromatin states and high expression.

The distributions of active histone modifications relative to gene expression suggested that not all highly expressed genes exhibited this canonical active-promoter profile. Indeed, Carbonic Anhydrase-2, a key coral skeleton-associated protein, was highly expressed (RPKM=4404) despite lacking strong H3K4me3 and H3K27ac peaks near the TSS (promoter log2(IP/input) score = −0.22, −0.18 respectively), while similarly expressed Elongation Factor 1-alpha (*EF1a,* RPKM=2654), exhibited strong active-mark promoter enrichment (promoter log2(IP/input) score = 0.98, 0.92 respectively; Figure 2E). This contrast suggested that some highly expressed coral genes may exhibit divergent promoter chromatin states.

### Expressed genes exhibit divergent promoter chromatin states and motif features

After identifying several highly expressed genes with divergent active-mark promoter enrichment, we next compared expressed genes by their promoter chromatin state. Genes with RPKM > 10 were divided into “High” and “Low” groups based on H3K4me3 and H3K27ac signal at promoters. The High group (n=7,418 genes) exhibited strong bimodal enrichment around the TSS, while the Low group (n=1,273) showed weaker, more diffuse promoter signal (Figure 3A,C). These differences were not explained by expression level, since these two groups had very similar mean and median RPKM values (High: 58.6 and 22.8; Low: 50.8 and 22.3, respectively).

**Figure 3.**
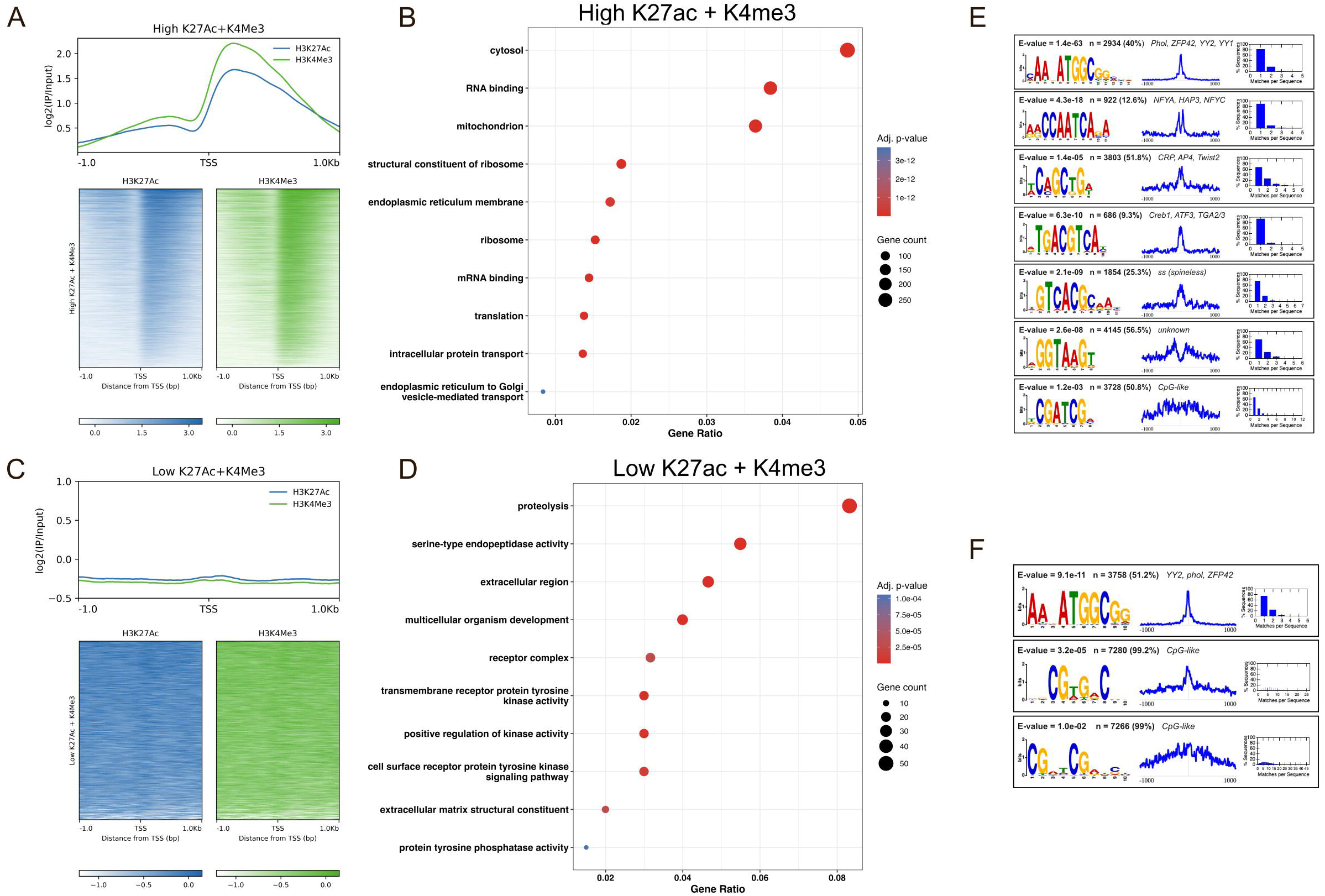
Expressed genes exhibit divergent promoter chromatin states and motif features. A) Log2(IP/input) H3K4me3 and H3K27ac signal across High active-mark promoters of expressed genes, defined as RPKM > 10 and mean promoter log2(IP/ input) > −0.1 for both marks (n= 7,418, mean RPKM= 58.6, median RPKM= 22.8). B) GO enrichment for High active-mark genes. C) H3K4me3 and H3K27ac signal across Low active-mark promoters of expressed genes, defined as RPKM > 10 and mean log2(IP/input) < −0.1 for both marks (n= 1,273, mean RPKM= 50.8, median RPKM= 22.3). D) GO enrichment analysis Low active-mark genes. E) Selected STREME-discovered motifs in High promoters (±1kb from TSS) against shuffled versions of the same sequences, identifying sequence features enriched relative to base composition alone. E-values, frequency, distribution, and TOMTOM matches are presented alongside motifs. F) Selected differential STREME motifs in High promoters, using Low active-mark promoters as the negative control set, identifying features that differ between the two groups.

To assess whether these patterns reflected technical artifacts, we evaluated three possibilities. First, competitive remapping of RNA-seq reads to a combined *P. damicornis*/Symbiodiniaceae reference genome reduced gene coverage to a similar degree in both promoter classes (median decrease: High, 14.2%; Low, 13.1%), indicating that Low-group expression was not disproportionately explained by misassigned symbiont reads. Second, promoter mappability was near-maximal for both groups, suggesting highly mappable unique promoter sequences (median 1.00 for both; mean 0.9995 for Low, 0.9998 for High). Third, to test whether transcription start site misannotation displaced genuine peaks outside the RefSeq-derived promoter window, we measured distances from each TSS to the nearest *macs2*-called H3K4me3 peak. Whereas 94% of High promoters lay within 1 kb of a peak, only 2% of Low promoters did, and Low and High promoters were equally distant from H3K4me3 peaks that were unassigned to any RefSeq annotated TSS (median 33-34 vs 31-32 kb; Mann–Whitney P = 0.54–0.58 across replicates; Table S3). Misannotated transcription start sites would place genuine H3K4me3 peaks near Low-group genes but outside the analysis window, where they would be unassigned to any RefSeq annotated TSS. Instead, unannotated peaks were no closer to Low- than to High-group promoters.

GO enrichment analysis revealed distinct functional profiles: High active mark genes were enriched for cytosol, mitochondria, and translation-related terms, while the Low active mark genes were enriched for proteolysis, extracellular receptors, and receptor tyrosine kinase activity (Figure 3B,D). We then used STREME to test whether promoter sequence features differed between groups. High-group promoters contained a conserved transcription-factor motif repertoire, concentrated within ±200 bp of the TSS (Figure 3E): A YY1-family motif (best TOMTOM match: *phol*, an invertebrate YY1 ortholog; present in 40% of promoters; E = 1.4 × 10^−63^), an NFY/CCAAT-box motif (NFYA/HAP3/NFYC; 12.6%; E = 4.3 × 10^−18^), an AP4/bHLH motif (TFAP4/Twist2; 51.8%; E = 1.4 × 10^−5^), and a CRE palindrome (CREB1/ATF3; 9.3%; E = 6.3 × 10^−10^). Additional motifs included a match to the bHLH-PAS factor *spineless* (25.3%; E = 2.1 × 10^−9^) and CpG-containing motifs without clear regulatory matches (Supplementary Data 1,2). Differential STREME analysis using Low promoters as the control background identified the same YY1-family motif enriched in High over Low promoters, present in 51% of High promoters but only 15% of Low promoters (E-value = 9.1 x 10^−11^; best TOMTOM match: YY2, *Phol*; Figure 3F). In addition, some CpG-containing motifs with no known TOMTOM matches were differentially enriched in High promoters, but these also existed in Low promoters (∼10% difference between groups; E-values <0.05; Figure 3F; Supplementary Data 3,4). In contrast, analysis of Low promoters against either scrambled or High-group backgrounds yielded few motifs with only one reaching significance (E= 3.0 x 10^−2^; Supplementary Data 5-8).

We also compared gene-body enrichment of the repressive modifications between groups. Low active-mark genes had higher gene-body H3K27me3 signal than High active-mark genes (Wilcoxon rank-sum, BH-corrected p<0.001; Figure S3). H3K9me3 showed less obvious separation between groups (Figure S3). Ultimately, High and Low active-mark promoters differed not only in histone modification state but also in their gene functions and frequency of these TSS-proximal sequence motifs.

### Repressive histone modifications H3K27me3 and H3K9me3 are associated with distinct genomic elements in the coral genome

To compare the two repressive marks, we ranked genes by gene-body H3K27me3 or H3K9me3 signal and analyzed the top 1,000 loci for each mark. H3K27me3 enriched genes were associated with neurotransmitter GPCR-related terms (Figure 4A), including adrenergic, dopamine, octopamine, and histamine receptor-like genes (Table S2). Conserved domain searches supported this pattern, identifying hundreds of proteins with 7TM domains characteristic of GPCRs (Figure 4B). Several neurotransmitter GPCRs were clustered together in the genome, with broad H3K27me3 enrichment and low transcriptional activity, suggesting epigenetic repression of these loci (Figure 4C).

**Figure 4.**
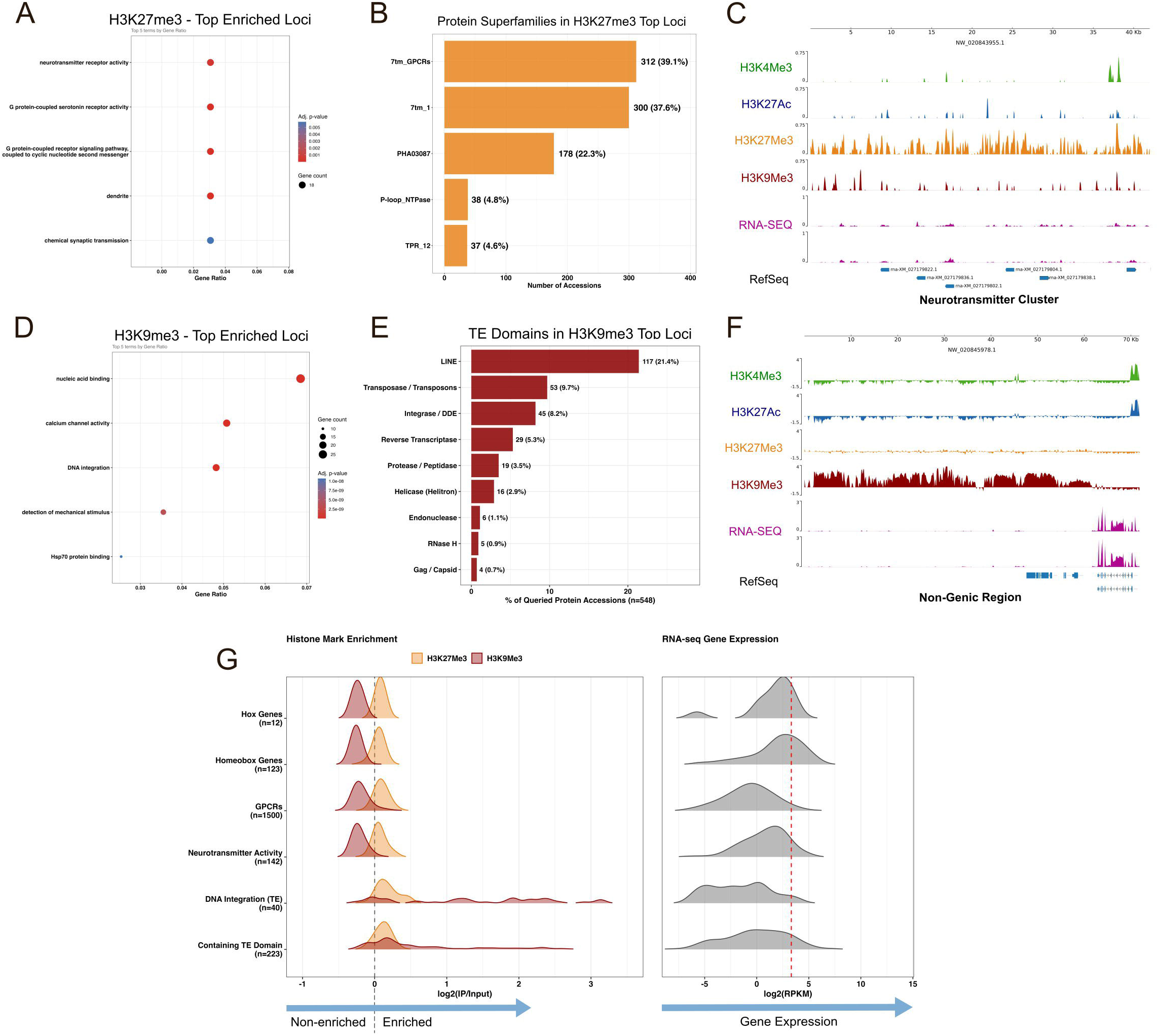
Repressive histone modifications H3K27me3 and H3K9me3 are associated with distinct genomic contexts in the coral genome. A) GO enrichment for the top 1,000 loci ranked by gene-body H3K27me3 signal. B) Most abundant conserved protein superfamilies among the top H3K27me3 loci. C) Log2(IP/ input) ChlP-seq and CPM-normalized RNA-seq tracks across a neurotransmitter receptor-like gene cluster. D) GO enrichment for the top 1000 loci as ranked by H3K9me3 signal across gene bodies. E) Frequencies of transposable element (TE) related protein domains among the top H3K9me3 loci. F) Log2(IP/input) ChlP-seq and CPM-normalized RNA-seq tracks across a large representative non-genie H3K9me3-enriched region. G) H3K27me3, H3K9me3, and RNA-seq expression distributions across selected repressed gene groups. Black dashed line indicates log2(IP/input) = 0; red dashed line indicates an RPKM > 10.

In contrast, the top 1,000 H3K9me3-enriched loci showed GO terms related to nucleic acid binding, calcium channel activity, and DNA integration (Figure 4D). Enrichment for calcium channel activity was driven by polycystin-like and TRPV-like proteins. Meanwhile, nucleic acid binding and DNA integration terms identified genes with endonuclease, integrase, and other transposable element (TE) related protein domains. Conserved protein domain searches further identified multiple TE-associated domains among the top H3K9me3 loci (Figure 4E). H3K9me3 signal was also observed across large non-genic regions, with negligible RNA-seq signal, consistent with broad constitutive heterochromatin domains (Figure 4F).

We then compared H3K27me3, H3K9me3 signal, and gene expression across relevant groups of genes. Hox-like genes, homeobox genes, GPCRs, and neurotransmitter-like GPCRs showed elevated H3K27me3 but comparatively low H3K9me3 signal (Figure 4G). In contrast, DNA integration genes and TE domain containing genes showed high signal for both repressive marks. Thus, similarly repressed gene groups displayed distinct repressive chromatin profiles, with H3K27me3 preferentially associated with neurotransmitter receptor-like GPCRs and developmental gene classes and H3K9me3 more strongly associated with TE-like and non-genic regions.

### Putative enhancer-like elements are proximal to genes in *P. damicornis*

In addition to promoters, we identified candidate enhancer-like elements as H3K27ac peaks outside the core TSS region (±500bp) with low H3K4me3 signal. Although H3K27ac peaks outside promoters were generally weaker than promoter-associated peaks, we identified 3,779 genic and 1,741 intergenic candidate enhancer-like regions. Both groups showed significantly higher H3K27ac than H3K4me3 signal across peak regions (Figure 5A-C), distinguishing them from characteristic active promoters (Figure 3).

**Figure 5.**
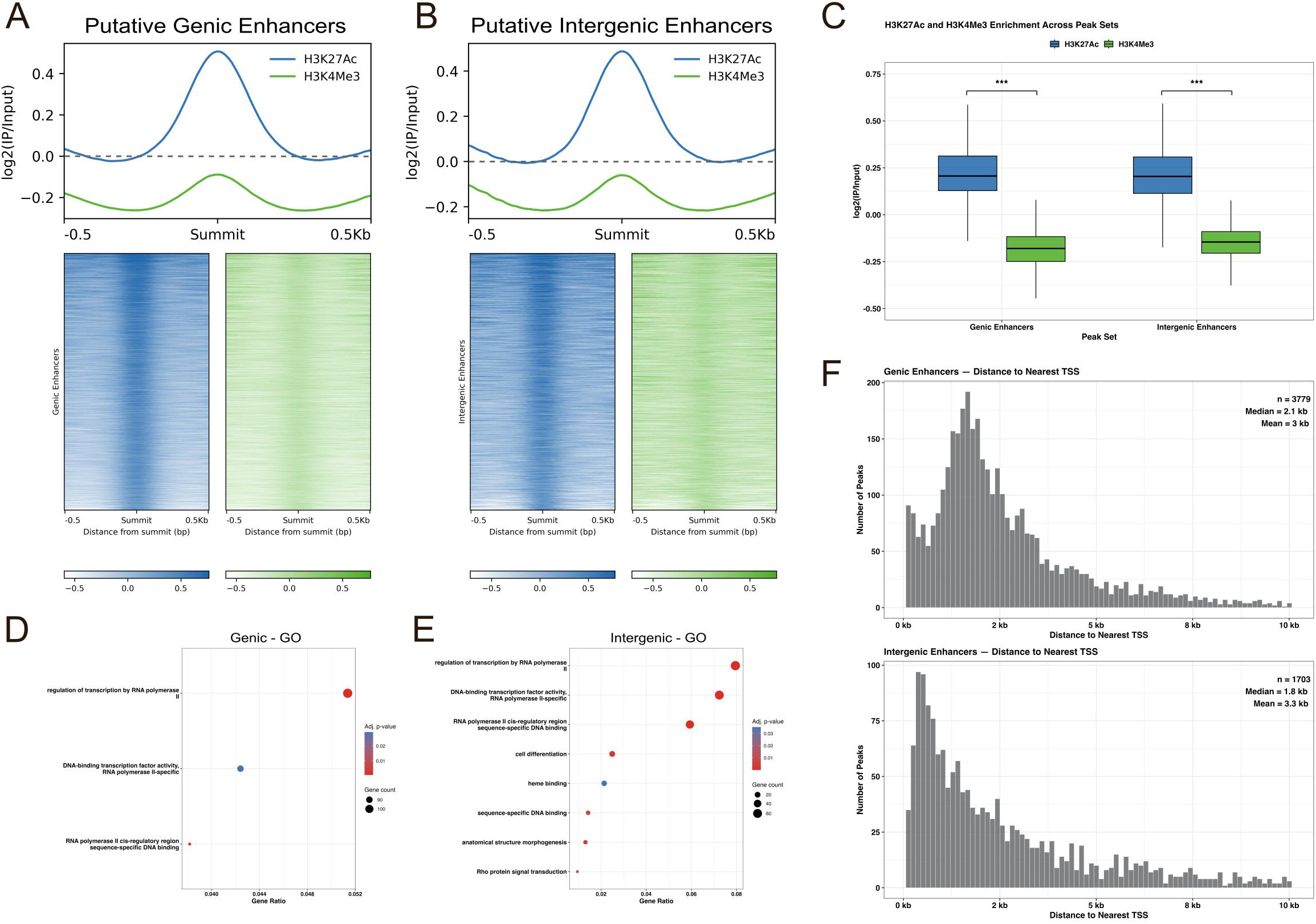
Putative enhancer-like elements are proximal to promoters in *P damicornis*. A,B) H3K27ac and H3K4me3 signal across putative genie enhancer-like regions (left, n= 3,779) and putative intergenic enhancer-like regions (right, n= 1,741). C) Mean log2(IP/input) signal for H3K27ac vs. H3K4me3 across candidate enhancer-like regions; significance was assessed by SH-corrected Wilcoxon tests. D,E) GO enrichment for genes annotated to putative genie and intergenic enhancer-like regions. F) Distance to nearest TSS for putative genie enhancers (top, mean= 3kb, median= 2.1kb) and intergenic enhancers (bottom, mean= 3.3kb, median= 1.8kb).

To infer potential functions of these regions, we assigned candidate enhancer-like peaks to their nearest TSS and performed GO enrichment analysis. Genes nearest to both genic and intergenic candidate enhancers were enriched for transcription factor-related terms (Figure 5D,E). These regions were also highly proximal to transcriptional start sites: Genic and intergenic candidates were, on average, 3.0 kb and 3.3 kb from the nearest TSS, respectively, with medians of 2.1 kb and 1.8 kb (Figure 5F).

Finally, STREME motif discovery across candidate enhancer-like elements identified motifs with variable patterns in signal and frequency and few matches to known transcription factors (Supplementary Data 9,10). Overall, these findings support the presence of H3K27ac-marked enhancer-like elements outside promoters in *P. damicornis*, many of which are located close to genes involved in transcriptional regulation. Further functional validation will be required to determine their regulatory activity.

## Discussion

This study establishes the first genome-wide maps of histone post-translational modifications in a reef-building coral, and shows that under baseline conditions, the *P. damicornis* genome contains conserved active and repressive chromatin states associated with transcriptional activity, functional gene classes, transposable element repression, and enhancer-like elements proximal to genes.

### A conserved chromatin logic is present in reef-building corals

Our results show that major histone modifications in *P. damicornis* follow conserved genomic distributions associated with transcriptional activity and repression. H3K4me3 and H3K27ac were enriched around transcriptional start sites and positively correlated with gene expression, whereas H3K27me3 and H3K9me3 showed broader enrichment patterns and were negatively associated with expression (Figures 1,2). This overall organization is consistent with genome-wide chromatin studies in the cnidarians *Aiptasia* (Nawaz et al., 2022) and *Nematostella* (Schwaiger et al., 2014), as well as in the sponge *Amphimedon* (Gaiti et al., 2017), supporting deep evolutionary conservation of active and repressive chromatin states across early-branching metazoans. Importantly, the coral epigenome was not simply divided into active and inactive regions: Histone modification enrichment varied across gene classes, scaffold gene density, and candidate regulatory elements, suggesting that these conserved chromatin states are deployed in biologically distinct genomic contexts in *P. damicornis*.

### Expressed genes in *P. damicornis* fall into two distinct promoter classes

Although H3K4me3 and H3K27ac ChIP signal in promoters was positively correlated with gene expression, this relationship broke down for a subset of genes. Among genes with RPKM > 10, we identified two groups with similar expression distributions but divergent promoter chromatin states. One carried the canonical bimodal H3K4me3/H3K27ac enrichment around the TSS, while a smaller group showed signal at or below background across the promoter window. Elongation Factor 1-alpha (*EF1a)* and Carbonic Anhydrase-2 illustrate this contrast: Both were highly expressed, but only EF1a showed the canonical H3K4me3/H3K27ac promoter signature.

We evaluated three technical possibilities that could account for this observation: 1) Competitive mapping against a combined host/Symbiodiniaceae reference excluded the possibility of misassigned symbiont reads. 2) Promoter mappability was near-maximal and indistinguishable between High and Low active-mark groups, ruling out poorly mappable sequences as an explanation. 3) Almost all Low promoters lacked H3K4me3 peaks within 1 kb of their TSS, and their equal distance to unassigned peaks relative to High promoters argued against TSS misannotation as the cause.

These two groups also differed functionally and in promoter sequence features. High active-mark genes were enriched for core cellular processes, including translation, mitochondrial activity, and cytosolic functions, and their promoters harbored a conserved metazoan transcription-factor repertoire concentrated within ±200 bp of the TSS: YY1-family, NFY/CCAAT-box, CRE/ATF, and AP4-bHLH motifs, together with CpG-containing motifs (Figure 3B,E,F). In contrast, Low active-mark genes were enriched for proteolysis, extracellular receptors, and kinase activity, and their promoters lacked this motif repertoire almost entirely (Figure 3D; Supplementary Data 5–8). These findings are consistent with promoter architectures in *Nematostella*, where YY1 motifs are enriched in constitutive promoters, while cell type-specific promoters lack these motifs (Elek et al., 2026). Moreover, YY1 motifs have been found at self-interacting domain boundaries in *Nematostella*, and this factor acts as a regulatory protein in promoter-enhancer interactions in vertebrates (Kim et al., 2025; Weintraub et al., 2017). Thus, our findings suggest that YY1 may play a similar regulatory role in the constitutive promoters of reef-building corals.

If the Low group corresponds to cell type-restricted promoters, its combination of high transcript abundance and low promoter signal is consistent with how ChIP-seq and RNA-seq scale in bulk tissue. Promoter ChIP signal is bounded by the number of marked promoter copies per nucleus and is therefore diluted roughly in proportion to the fraction of cells carrying the mark, whereas transcript abundance lacks these limitations: A gene transcribed intensely in a minority of cells could be interpreted as moderate expression across all cell types in bulk RNA-seq libraries. Increased gene-body H3K27me3 signal in the Low group is also consistent with this interpretation, suggesting repression in most cells where these genes would be inactive (Figure S3). Therefore, the difference between these promoter groups may lie in the cell-type specificity of their expression.

However, dilution is unlikely to fully explain these differences in promoter classes. For example, Carbonic Anhydrase-2 is expressed in both calicoblasts and symbiocytes (Han et al., 2025), which constitute a substantial fraction of coral tissue, so its weak promoter signal cannot readily be attributed to expression in a rare cell type. Therefore, other regulatory configurations may also contribute to these findings. First, nucleosome-depleted promoters would yield low ChIP signal for active marks irrespective of modification state. Second, the four modifications profiled here are all on histone H3 and represent a small subset of the coral chromatin landscape. Diverse modifications and sequence variants on histones H4, H2A, and H2B, as well as non-canonical histone processing have been reported in corals (Fuller et al., 2025b; Rodriguez-Casariego et al., 2018; Roquis et al., 2022). Last, transcription of these genes may also depend on DNA methylation, which is coupled to histone modification state in *Aiptasia* (Li et al., 2018; Nawaz et al., 2022). Ultimately, distinguishing among these possibilities will require chromatin and transcriptional profiling at single-cell resolution.

### Repressive histone modifications define distinct inactive chromatin states

H3K27me3 and H3K9me3 were both negatively correlated with gene expression in *P. damicornis*, but their genomic distributions suggest distinct repressive roles in the genome. H3K27me3 was enriched across broad genic regions with low transcriptional activity, particularly over neurotransmitter receptor-like GPCRs, including adrenergic, dopamine, octopamine, and histamine receptor-like genes, as well as homeobox/Hox-like genes (Figure 4A,B,C). This pattern is consistent with the canonical role of broad H3K27me3 domains in transcriptional silencing (Young et al., 2011) and with H3K27me3-associated repression reported in *Aiptasia* (Nawaz et al., 2022) and the sponge *Amphimedon* (Gaiti et al., 2017), supporting a conserved role for this mark in cnidarian gene regulation. In corals, these H3K27me3-marked loci may represent genes, such as GPCRs, that are repressed in most adult cells but are active in distinct cell types, developmental stages, or physiological contexts (Han et al., 2025; Levy et al., 2021; Moeller et al., 2019).

In contrast, H3K9me3 was most prominent over TE-like loci and broad non-genic regions with low transcriptional activity, consistent with a constitutive heterochromatin-like state (Figure 4D,E,F). This pattern resembles the conserved association of H3K9 trimethylation with transcriptionally silent repetitive elements and heterochromatin in other eukaryotes (Padeken et al., 2022), suggesting that H3K9me3 may similarly contribute to repression of repeat-associated or mobile-element-like regions in the coral genome. TE-like loci also showed enrichment of H3K27me3 (Figure 4G), suggesting that these two repressive marks may overlap at these genomic elements, even though broader genomic patterns of these marks suggest distinct primary roles. Thus, coral repressive chromatin appears to be functionally partitioned by H3K27me3 at regulated gene loci and by H3K9me3, at non-genic heterochromatin-like regions, with partial overlap at silenced TE-like loci.

### H3K27ac-marked regions outside core promoters lie within a compact genomic context

We also identified H3K27ac-enriched, H3K4me3-low regions outside the core TSS window that may represent candidate enhancer-like elements in *P. damicornis*. These regions had weaker H3K27ac peaks than active promoters, but were significantly enriched for H3K27ac relative to H3K4me3, supporting their distinction from canonical promoter-associated active chromatin (Figure 5A,B,C). Because these elements were identified by chromatin state rather than functional validation, we refer to them as putative enhancer-like regions rather than confirmed enhancers. A notable feature of these regions was their close proximity to genes. Both genic and intergenic candidates were located close to transcriptional start sites, with median distances of approximately 2.1 kb and 1.8 kb, respectively (Figure 5F). These short distances are consistent with the compact spacing of annotated genes in the *P. damicornis* genome, where mean and median intergenic distances are 3.1 kb and 1.4 kb, respectively, and are much smaller than those found in more complex animals, such as humans (Gherman et al., 2009). These observations suggest that candidate cis-regulatory elements in *P. damicornis* may be positioned within small intergenic gaps, highly proximal to target promoters.

This compact organization is consistent with what is known of cis-regulatory architecture in other cnidarians. Chromatin profiling in the anemone *Nematostella* (Schwaiger et al., 2014) and single-cell chromatin accessibility profiling has since resolved over 100,000 candidate cis-regulatory elements across cell types, assigning peaks to genes within a 10 kb window (Elek et al., 2026). In *Nematostella* and *Scolanthus*, cis-regulatory sequences tend to lie close to their target promoters, consistent with the compact nature of cnidarian genomes (Zimmermann et al., 2023). The median distances we observed in *P. damicornis* (1.8–2.1 kb) fall well within these ranges. Without Hi-C or functional validation in *P. damicornis*, we view this as evolutionary context rather than a direct description of coral genome organization. Our findings indicate that the *P. damicornis* genome contains H3K27ac-marked regions outside core promoters, whose nearest genes are enriched for transcription factor-related functions (Figure 5D,E).

### Considerations

The ChIP-seq profiles obtained here were reproducible and biologically interpretable across multiple histone modifications. Antibody validation, replicate correlations, separation from input controls, H3K4me3/H3K27ac in active promoters, and broad H3K27me3/H3K9me3 signal over lowly expressed or non-genic regions all support the successful recovery of positional information of histone modification enrichment in reef-building coral nuclei. We expect that careful tissue dissociation, fixation, chromatin extraction, and sonication were particularly important, as these steps determine whether chromatin is preserved, accessible to antibodies, and fragmented at appropriate sizes for immunoprecipitation. Importantly, a single genotype cannot capture inter-individual variation in chromatin state, and that generalization across genotypes awaits future work.

Bulk ChIP-seq and RNA-seq provide a tissue-level view of chromatin state and transcriptomic output, effectively averaging signal across many cell types present in coral tissue. Because of this, cell-type specific regulatory elements and expression patterns can be diluted (see discussion of Promoter Classes). Thus, while bulk ChIP-seq cannot resolve cell-type-specific chromatin states, it remains useful for identifying tissue-level chromatin features and for identifying epigenetic shifts under stress conditions, which typically elicit strong tissue-level transcriptional responses.

### Outlook: Chromatin mapping as a framework for coral environmental response

By establishing genome-wide histone modification maps in a reef-building coral, this study provides a baseline framework for connecting coral genome organization with transcriptional regulation. Coral stress responses are often studied through changes in gene expression, but transcript abundance represents the downstream outcome of regulatory processes acting at promoters, gene bodies, repressive chromatin domains, and regulatory regions outside promoters. Mapping histone modifications therefore adds a regulatory layer for distinguishing active, repressed, and enhancer-like chromatin signatures in coral tissue. These maps also identify coral loci with known bulk chromatin signatures that can serve as positive and negative validation controls in targeted assays such as ChIP-qPCR. More broadly, this work addresses a foundational gap in coral epigenomics by mapping histone modification landscapes in a reef-building coral and providing a foundation for asking how coral chromatin states are remodeled across environmental, developmental, and symbiotic contexts.

An important next step will be to integrate ChIP-seq with other genomic techniques, such as ATAC-seq, Hi-C, DNA methylation profiling, and RNA-seq. Combining these approaches in reef-building corals could reveal how chromatin accessibility, 3D genome organization, and histone modification states jointly shape transcriptional responses to stress. These multi-omic approaches will be especially informative at the single-cell level, helping to identify cell-type specific changes that contribute to bulk regulatory and transcriptional signals.

## Methods

### Coral Husbandry and Experimental Setup

*Pocillopora damicornis* (single genotype, originally obtained from Battlecorals in Wisconsin, USA) was maintained in 100-liter aquaria at a temperature of 25C, 35 ppt salinity, 7-8 dKH carbonate hardness, 400-450ppm Calcium, ∼1350ppm Magnesium. Water chemistry was maintained through weekly 20% water changes with artificial seawater and by scheduled dosing of calcium chloride, sodium carbonate, and magnesium chloride solutions. Variable flow in the aquaria was attained using two RPM Wavemakers (9,800 l/h capacity each; ReefBreeders). Photon V2-Pro LED lights (ReefBreeders) were mounted 35cm above coral fragments and ran on a 12-hour ON (ramping) and 12-hour OFF schedule to mimic natural reef lighting. This fixture uses a combination of 36 individually dimmable LEDs across six spectral channels: Deep red (660 nm), green (520 nm), royal blue (450 nm), white (5500 K), cool blue (480 nm), and violet (420 nm), totaling 144 W of nominal LED capacity.

To control for genotypic variation, all material was derived from a single parent colony, which was fragmented into pieces ∼5cm in height, and fragments were allowed to recover and heal for 6 weeks prior to sampling. Two fragments, BR1 (Rep 1) and BR2 (Rep 2), maintained separately in the same system, were harvested independently and processed as isogenic replicates; each yielded a pooled chromatin preparation that was divided across four immunoprecipitations. For H3K27ac, chromatin from BR1 was immunoprecipitated twice BR1 (Rep 1) and BR1B (Rep 1B) to assess technical reproducibility.

### Histone H3 Protein Sequence Alignment

To investigate histone protein conservation amongst species, protein sequences from human, mouse, *Aiptasia* anemones, and the reef-building corals *Pocillopora damicornis*, *Stylophora pistillata, Montipora capricornis,* and *Acropora millepora* were aligned using COBALT in NCBI. Alignments were visualized using the Python package *pymsaviz* (Figure S1A).

### Western Blots with ChIP Antibodies

To extract total proteins, coral tissue was sampled using bone cutters and flash frozen in LN2. Coral tissue was cut into small pieces under LN2, then placed into TRIzol reagent (Invitrogen). Tissue was homogenized using a handheld motorized tissue homogenizer (DWK Life Sciences), and protein extraction was performed according to the manufacturer’s protocol. Protein yield was quantified using a Bradford Assay on a Nanodrop Spectrophotometer (ThermoFisher). In Bis-Tris gels, 30µg of total protein in 1X loading and reducing buffer (Invitrogen) were loaded into each lane. Blots were stained with rabbit primary antibodies (anti-H3K27ac [Abcam, ab4729], anti-H3K27me3 [Diagenode, C15410069], anti-H3K4me3 [Abcam, ab8580], or anti-H3K9me3 [Abcam, ab176916]) and HRP-conjugated Goat anti-Rabbit secondary antibodies using the iBind Flex kit (Invitrogen). Western blot fluorescence (Figure S1B) was visualized using the Pierce Chemiluminescent kit (ThermoFisher).

### Coral Tissue Fixation and Dissociation for ChIP-seq

Coral tissue was trimmed from two isogenic replicates using bone-cutters, briefly rinsed in 0.6M NaCl-supplemented PBS (0.6M NaCl, 1x PBS), and placed in 1% paraformaldehyde Fixative Buffer (0.6M NaCl, 1x PBS, 1% paraformaldehyde). Fragments were incubated on a rocker for 15 minutes. To quench the fixation, glycine was added to achieve 0.125M concentration, and samples incubated for 5 minutes on a rocker. Samples were rinsed twice with ice-cold 1x PBS and then submerged in ice-cold EDTA Storage Buffer (50 mM Tris-HCl [pH 8], 10 mM EDTA, 0.5 mM EGTA, 140 mM NaCl) with 1X Protease Inhibitor (Thermo Scientific) prior to weighing. Wet tissue sample masses were measured, and tissue was returned to the EDTA Storage Buffer. To dissociate the tissue, 0.2µm-filtered pressurized air was channeled through a 200µl pipette tip and aimed at the surface of the coral tissue. The tissue was sprayed directly into ice-cold EDTA Storage Buffer with 1X Protease Inhibitor (Thermo Scientific). The resulting tissue slurry was centrifuged at 600 x g for 15 minutes to pellet the cells. After removal of the supernatant, cell pellets were snap frozen in liquid nitrogen and stored at −80C.

### Coral Chromatin Extraction and Sonication

Frozen cell pellets from the two original replicates were thawed on ice and resuspended in Lysis Buffer (50 mM Tris-HCl [pH 8], 10 mM EDTA, 0.5 mM EGTA, 140 mM NaCl, 1% SDS) with 1X Protease Inhibitor (Thermo Scientific). Cell suspensions were homogenized using a Dounce homogenizer and left to incubate for an additional 15 minutes on ice. Samples were sonicated using the BioRuptor Pico (Diagenode) for 8 cycles of 15sec ON:30sec OFF to target an optimal chromatin fragmentation of 150-300bp (Figure S1C). After sonication, samples were centrifuged at 21,130 x g at RT for 10 minutes. The resulting solubilized chromatin was diluted with ChIP Dilution Buffer (16.7 mM Tris-HCl [pH = 8.1], 1.2 mM EDTA, 167 mM NaCl, 0.01% SDS, 1.1% Triton X-100) with 1X Protease Inhibitor (Thermo Scientific) at a ratio of 5mL dilution buffer:1mL solubilized chromatin. At this time, aliquots of the diluted chromatin were removed to serve as input controls for the immunoprecipitations and stored at 4C until decrosslinking.

### Immunoprecipitation

Pools of diluted chromatin from each original replicate were split into aliquots for four immunoprecipitations, each containing around 25µg total DNA extracted from ∼900mg of tissue (tissue + skeleton wet weight). 2µg of anti-H3K27ac (Abcam, ab4729), anti-H3K27me3 (Diagenode, C15410069), anti-H3K4me3 (Abcam, ab8580), or anti-H3K9me3 (Abcam, ab176916) were added to respective immunoprecipitation reactions. Samples were incubated overnight on a rotator at 4C. The following day, Protein A Dynabeads (Invitrogen) were added for immunoprecipitation, and tubes were incubated for 2 hours on a rotator at 4C. Beads were pelleted out of solution and washed 2x with of ice-cold Low-Salt Immune Complex Wash Buffer (20mM Tris-HCl [pH=8.1], 2mM EDTA, 150mM NaCl, 0.10% SDS, 1% Triton X-100), 2x with ice-cold High-Salt Immune Complex Wash Buffer (10 mM Tris-HCl [pH 8.1], 1 mM EDTA, 0.25 M LiCl, 1% Deoxycholic Acid, 1% NP-40), and 2x with ice-cold 1X TE Buffer, pH 8.0. Beads were resuspended in DTT Elution Buffer (10 mM Tris-HCl [pH 8.1], 1 mM EDTA, 150 mM NaCl, 1% SDS, 5 mM DTT) and incubated at 65C. After pelleting the beads with a magnet, the chromatin-containing supernatant was recovered. At this point, NaCl, SDS, and DTT were added to the 500µl input controls (WCEs). All samples were incubated overnight at 65C to de-crosslink proteins from DNA.

### DNA Purification

De-crosslinked samples were treated with proteinase K and incubated at 55C for 2 hours. DNA was extracted using phenol-chloroform, and DNA was precipitated overnight at −20C in 100% ethanol, supplemented with sodium acetate and glycogen for DNA pellet visualization. After overnight incubation, samples were centrifuged at 21,130 x g for 30 minutes at 4C. DNA pellets were rinsed with 70% ethanol, and the samples were centrifuged at 21,130 x g for another 15 minutes. The ethanol supernatant was removed, and pellets were dried in a fume hood until glossy. DNA pellets were eluted in a 1:50 dilution of RNAse A in 1X TE buffer, pH 8.0. Eluted DNA was incubated at 37C for 30 minutes to remove RNA contamination. DNA yields were quantified using Qubit HS dsDNA assay kit (Invitrogen). Input control and immunoprecipitated DNA samples were stored at −20C until library preparation.

### ChIP DNA Library Prep and Sequencing

Libraries were prepped using the NEBNext Ultra II DNA Library Prep Kit for Illumina (New England Biolabs) and indexed with NEBNext Multiplex Unique Dual Index Oligos (New England Biolabs), following the manufacturers protocols. Library yield was quantified using Qubit HS dsDNA assay kit (Invitrogen) and fragment size distribution was checked on a HS DNA bioanalyzer chip (Agilent) prior to pooling libraries for sequencing. Samples were sequenced in multiplex on a 1000M PE150 NovaSeq X Plus flow cell (Illumina) through NUSeq Core (Northwestern Center for Genetic Medicine).

### ChIP-seq Quality Control and Mapping

An average of 36 million 150bp read pairs were obtained for each ChIP-seq library (Table S4). Sequencing quality was verified using *FastQC*. Adapters and poor quality reads were detected and removed using *fastp* (Chen et al., 2018). Paired-end reads were mapped using *bwa mem* (Li & Durbin, 2009) to a combined reference (RefSeq) genome containing the *Pocillopora damicornis* (GCF_003704095.1) genome and two algal symbiont genomes: *Durusdinium trenchii* (GCA_963970005.1) and *Cladocopium goreaui* (GCA_947184155.2). Mapping quality was assessed using *samtools*. Reads that mapped to the coral genome with a mapping score (MapQ) greater than 30 were processed using *Picard* to remove PCR and optical duplicates.

### RNA Extraction

Coral tissue from each replicate was sampled using bone cutters, rinsed in 0.6M NaCl-supplemented PBS, and flash frozen in LN2. Frozen tissue was stored at −80C until RNA extraction. To extract total RNA, samples were cut under liquid nitrogen, then homogenized in DNA/RNA Shield (ZYMO) using a handheld motorized tissue homogenizer (DWK Life Sciences). The homogenate was then processed using the ZYMO Quick-DNA/RNA Miniprep Plus Kit. After elution, all extractions were passed through OneStep PCR Inhibitor Removal columns (ZYMO). DNA and RNA yields were quantified using a Qubit Fluorometer (Invitrogen). RNA extracts were analyzed for RNA integrity using an Agilent TapeStation prior to RNA-seq library preparation.

### RNA-seq Library Prep and Sequencing

Using the NEBNext Poly(A) mRNA Magnetic Isolation Module (New England Biolabs), RNA was isolated from 200ng of total RNA extracted from each replicate (n=2). Isolated poly(A)-RNA was fragmented to a target length of 200bp and reverse-transcribed into cDNA libraries using the NEBNext UltraExpress RNA Library Kit (New England Biolabs). After adapter ligation, NEBNext Multiplex Unique Dual Index Oligos (New England Biolabs) were used for library indexing. RNA-seq library yields were quantified using Qubit (Invitrogen) and the size distribution was determined using an Agilent Bioanalyzer prior to sequencing. RNA-seq libraries were sequenced in multiplex using NovaSeq X Plus (Illumina) 50bp paired-end sequencing through NUSeq Core (Northwestern Center for Genetic Medicine).

### RNA-seq Quality Control and Mapping

Adapters and poor-quality reads were detected and removed using *fastp* from 73.8 million raw 50bp read pairs for replicate 1 and 57.2 million read pairs for replicate 2 (Table S5). Sequencing quality was analyzed using *FastQC*. Paired-end 50bp reads were mapped using *STAR* (Dobin et al., 2013) to the *Pocillopora damicornis* (GCF_003704095.1) genome. The following arguments were used in *STAR* as recommended by ENCODE:

--outFilterMultimapNmax 20,
--outFilterMismatchNoverReadLmax 0.04,
--alignSJoverhangMin 8,
--alignSJDBoverhangMin 1,
--alignIntronMin 20,
--alignIntronMax 1000000,
--alignMatesGapMax 1000000,
--outFilterIntronMotifs RemoveNoncanonical.

Direct mapping to RefSeq annotations was used to obtain read counts for RefSeq genes. Raw counts were normalized by library size by calculating size factors in *DESeq2* and by gene length using RPKM normalization in *edgeR* (Love et al., 2014; Robinson et al., 2010). Mitochondrial and tRNA transcripts were removed. Reads were also mapped to generate stranded and unstranded CPM-normalized BigWig files for RNA-seq data.

### Genomic Distributions of Histone Modifications

BigWig files were generated using *deepTools* BamCoverage and counts per million (CPM) normalization (Ramírez et al., 2014). Average BigWig tracks were created for each mark across both replicates using *deepTools* bigwigAverage, which were used to visualize the ChIP signal of each modification and input controls across regions of interest (such as RefSeq gene bodies) using *deepTools* plotProfile and plotHeatmap. To visualize ChIP signal at specific loci, CPM-normalized BigWig tracks were plotted in *pyGenomeTracks* (Lopez-Delisle et al., 2021). Using *deepTools* bamCompare, log2(IP/input) BigWig tracks were generated for each histone modification, averaged across replicates.

Correlation between histone marks was determined by comparing CPM-normalized read counts in 1000bp bins throughout the entire genome using *deepTools* multiBigwigSummary. A greylist of artifact peak regions that appeared in input control libraries was created using the *GreyListChIP* package in R. The greylist was applied to the called peak files for the immunoprecipitation libraries, and any overlapping peaks were discarded using *bedtools* (Quinlan & Hall, 2010). Spearman correlation coefficients were determined for all libraries and visualized in a heatmap using *deepTools* plotCorrelation, after applying the input control greylist. Mean log2(IP/input) scores for each histone modification across promoters (±1kb from TSS) of all RefSeq genes were computed using bigwigCompare and multiBigwigSummary. Correlation scatterplots and spearman coefficients for each histone mark pairwise comparison was visualized using plotCorrelation, after applying the greylist.

To visualize how histone modification enrichment related to gene density in scaffolds, the genic density of each scaffold was computed by dividing the overlapped length of all genes in the scaffold by the total length of the scaffold. Shorter and unreliable scaffolds (<5000bp) were removed. The mean log2(IP/input) score across each scaffold was computed using *deepTools* multiBigwigSummary. Scaffolds containing zero genes were labeled “non-genic,” and the remaining genic scaffolds were split into four quartiles by gene density. Unpaired, Benjamini Hochberg (BH)-corrected, Wilcoxon rank-sum tests were performed for the non-genic group vs. all genic groups (n=4 tests). Effect size was determined from the W test statistic.

Narrow peaks in H3K4me3 and H3K27ac were called in *macs2* using a q-value cutoff of 0.05, an approximate effective genome size of 2.25e8 (all non-N bases in the assembly), and the --BAMPE argument to account for PE150bp reads. Peaks were annotated to genomic features using *ChIPseeker* in R (Yu et al., 2015), and frequencies for different genomic features were visualized using bar plots.

### Histone Modifications and Gene Expression

Bed files for the promoters (±1kb from TSS) and gene bodies (TSS-TES) for all RefSeq genes were inputted into *deepTools* MultiBigWigSummary to obtain mean log2(IP/input) scores of histone modifications across each region. RefSeq transcripts not detected by RNA-seq (n=1546) were denoted as “undetected”, and the remaining genes with RPKM > 0 were partitioned into quartiles (n=5297 each) by their RPKM expression. Log2(IP/input) signal across RefSeq genes (+2kb from TSS/TES) was visualized using *deepTools* plotProfile.

Using all detected RefSeq genes, the Spearman correlation coefficient between log2(IP/input) scores in promoters (H3K4me3, H3K27ac) or gene bodies (H3K27me3, H3K9me3) and log2(RPKM) expression was determined. Scatter plots and linear fits were visualized using *ggplot2* in R. Selected functional groups of genes were classified using Gene Ontology terms and keywords to visualize their chromatin and gene expression states using ridgeline plots (*ggridges* in R). Two highly expressed genes of interest with divergent chromatin states, Elongation Factor 1-alpha (RPKM=2654) and Carbonic Anhydrase-2 (RPKM=4404), were visualized in *pygenometracks* using ChIP-seq and RNA-seq BigWig tracks.

### Gene Expression and Chromatin State

Transcripts with RPKM > 10 were selected as genes with strong expression (n=8,995). Using a log2(IP/input) promoter score cutoff of −0.1 for H3K4me3 and H3K27ac, these genes were partitioned into “High” (n=7,418, mean RPKM=58.6, median RPKM=22.8) and “Low” (n=1,273, mean RPKM=50.8, median RPKM=22.3) groups by chromatin state in their promoters. Promoters with one mark above the cutoff and the other below (n=304) were excluded. H3K4me3 and H3K27ac log2(IP/input) signal was visualized using *deepTools* plotProfile and plotHeatmap across promoters (±1kb from TSS). Gene ontology (GO) enrichment analysis was performed for each group using *ClusterProfiler* and Benjamini Hochberg (BH) correction (minGSSize=10, maxGSSize=500). A background of all expressed genes (RPKM > 0) with computed ChIP promoter scores was used. To ensure that RNA-seq signal was not contaminated by algae-originating reads, the RNA-seq was remapped to a combined *Pocillopora damicornis, Durusdinium trenchii,* and *Cladocopium goreaui* reference genome using the same *STAR* methods as above.

To ensure that the absence of activating mark enrichment in Low promoters were not inaccuracies from the existing RefSeq TSS annotation, we performed a sensitivity analysis: The genomic distance from each TSS in Low or High promoters to the most proximal H3K4me3 peak was measured using *bedtools* closest. This was performed against 1) all *macs2*-called peaks, and 2) the subset of H3K4me3 peaks not found on existing RefSeq TSS, which may belong to misannotated TSSs. Distance distributions were calculated, and the fraction of High and Low promoters that lay within 1 kb, 2 kb, and 5 kb distances from H3K4me3 peaks (Table S3).

STREME and TOMTOM were used to identify enriched motifs in these two groups. First, STREME was performed independently for each group using a randomly shuffled version of the same group’s sequences as the negative control set. Differential STREME was then performed for each group, using the opposite group as the negative control set to identify motifs unique to each group. TOMTOM was performed against the JASPAR 2026 CORE non-redundant database (Ovek Baydar et al., 2026). When isolating FASTA sequences within identified promoter groups, a small number of promoter sequences that occurred at the edges of scaffolds (where a full ±1kb from TSS was not possible) were removed from the analysis (High: n=80, 1.1%; Low: n=136, 10.7%).

### Analysis of Repressive Histone Modifications H3K27me3 and H3K9me3

All RefSeq genes were ranked by log2(IP/input) scores for H3K27me3 and H3K9me3 in their gene bodies. The top 1000 loci for each modification were selected after removal of potential false positives using the input control greylist. GO enrichment analysis was performed for each group using *ClusterProfiler* and Benjamini Hochberg (BH) correction (minGSSize=10, maxGSSize=500), using a background of all RefSeq genes with computed gene body ChIP scores. Loci with no GO annotation terms were excluded from analyses. Protein accessions for the top 1000 loci in each group were queried against the NCBI conserved domain database to identify protein domains and superfamilies within each locus. Loci without protein accessions were excluded from analysis. Domains associated with transposable elements (TEs) were identified using TE-related keywords, and their frequency was recorded within the top 1000 loci for each histone modification. The unique set of combined genes from both groups that contained these TE protein domains were labeled “Containing TE Domain.” Examples of H3K27me3 and H3K9me3 broad repressive chromatin states were visualized using *pygenometracks* using log2(IP/input) BigWig tracks. The chromatin state and gene expression of repressed groups of genes and TE-like loci were represented using a ridgeline plot.

### Investigation of Putative Enhancers

Peaks in H3K4me3 and H3K27ac were called in *macs2* using a q-value cutoff of 0.2, an approximate effective genome size of 2.25e8 (all non-N bases in the assembly), and --BAMPE for 150bp PE reads. A higher q-value cutoff of 0.2 was used in this case to generate a wider list of candidate enhancers. H3K27ac peaks were filtered to remove those overlapping with peaks in H3K4me3 using *bedtools* intersect. Additional filtering using a mean H3K4me3 log2(IP/input) score < 0 across the peak regions was also used. Peaks were recentered around their summits using the R package *DiffBind* and raw deduplicated BAM files. Only consensus peaks across all three H3K27ac replicates were kept. Putative enhancer peaks were split into “genic” (n=3779) and “non-genic” (n=1741) groups using *bedtools* intersect against RefSeq gene bodies. Signal across these regions was visualized using *deepTools* plotProfile and plotHeatmap. Differences between H3K27ac and H3K4me3 log2(IP/input) score distributions across these regions were analyzed using paired Wilcoxon signed-rank tests and BH correction for n=2 tests. Putative-enhancers were annotated to their nearest genes with *ChIPseeker*, and GO enrichment analysis was performed on genic and intergenic regions separately, using *ClusterProfiler,* BH correction (minGSSize=10, maxGSSize=500), and a background of all RefSeq genes. STREME and TOMTOM were used to identify conserved motif patterns in these putative enhancer regions, using the JASPAR 2026 CORE non-redundant database (Supplementary Data 9,10). In addition, the distance between enhancers and the nearest TSS was computed for peaks within each group, and distance distributions were visualized using histograms. When calculating genomic distances from the nearest TSS, a small number of enhancer-like regions that occurred on non-genic scaffolds (where a genomic distance value was not possible) were removed (n=38, 2% for intergenic regions; n=0, 0% for genic regions).

## Supporting information

Supplementary Data 1 - STREME High Promoters (scrambled background)

Supplementary Data 2 - TOMTOM High Promoters (scrambled background)

Supplementary Data 3 - STREME High Promoters (High vs. Low)

Supplementary Data 4 - TOMTOM High Promoters (High vs. Low)

Supplementary Data 5 - STREME Low Promoters (scrambled background)

Supplementary Data 6 - TOMTOM Low Promoters (scrambled background)

Supplementary Data 7 - STREME Low Promoters (Low vs. High)

Supplementary Data 8 - TOMTOM Low Promoters (Low vs. High)

Supplementary Data 9 - STREME Putative Enhancers

Supplementary Data 10 - TOMTOM Putative Enhancers

## Acknowledgements

We would like to thank the members of the Adli, Backman, and Marcelino labs, in particular Dr. Cody L. Dunton, Brittany Johnson, and Yasemin Tekin for many fruitful conversations. Thank you also to Anne Warfel for assistance with coral husbandry. This work was kindly supported by the Northwestern University NUSeq Core Facility, and in part through the computational resources and staff contributions provided by the Genomics Compute Cluster, which is jointly supported by the Feinberg School of Medicine, the Center for Genetic Medicine, Feinberg’s Department of Biochemistry and Molecular Genetics, the Office of the Provost, the Office for Research, the Weinberg College of Arts and Sciences, and Northwestern Information Technology. The Genomics Compute Cluster is part of Quest, Northwestern University’s high-performance computing facility, with the purpose of advancing research in genomics. This work was supported by NSF Grant CBET-2427519, an award from the Northwestern University Center for Physical Genomics and Engineering, and awards from the Ubben Program for Climate and Carbon Science and the Resnick Family Social Impact Fund at the Paula M. Trienens Institute for Sustainability and Energy, Northwestern University. GW was also supported by the Murphy Scholars Program of the Robert R. McCormick School of Engineering and Applied Science at Northwestern University. The graphical abstract was created in BioRender. Sadalla, L. https://biorender.com/hb2ikn7.

## Author Contributions

**GW:** Conceptualization, Methodology, Investigation, Formal Analysis, Writing - Original Draft. **SLD:** Methodology, Investigation; Validation. **LS:** Investigation, Analysis, Validation. **TV, FSP, HO:** Methodology. **VB:** Resources, Conceptualization. **MA:** Resources, Conceptualization, Supervision. **LAM:** Resources, Conceptualization, Supervision. **All:** Writing - Review and Editing.

## Data Availability

ChIP-seq and RNA-seq raw sequencing data will be available in the NCBI Sequence Read Archive (SRA) under BioProject PRJNA1515428, after the manuscript is accepted for publication.

---

**Figure S1.**
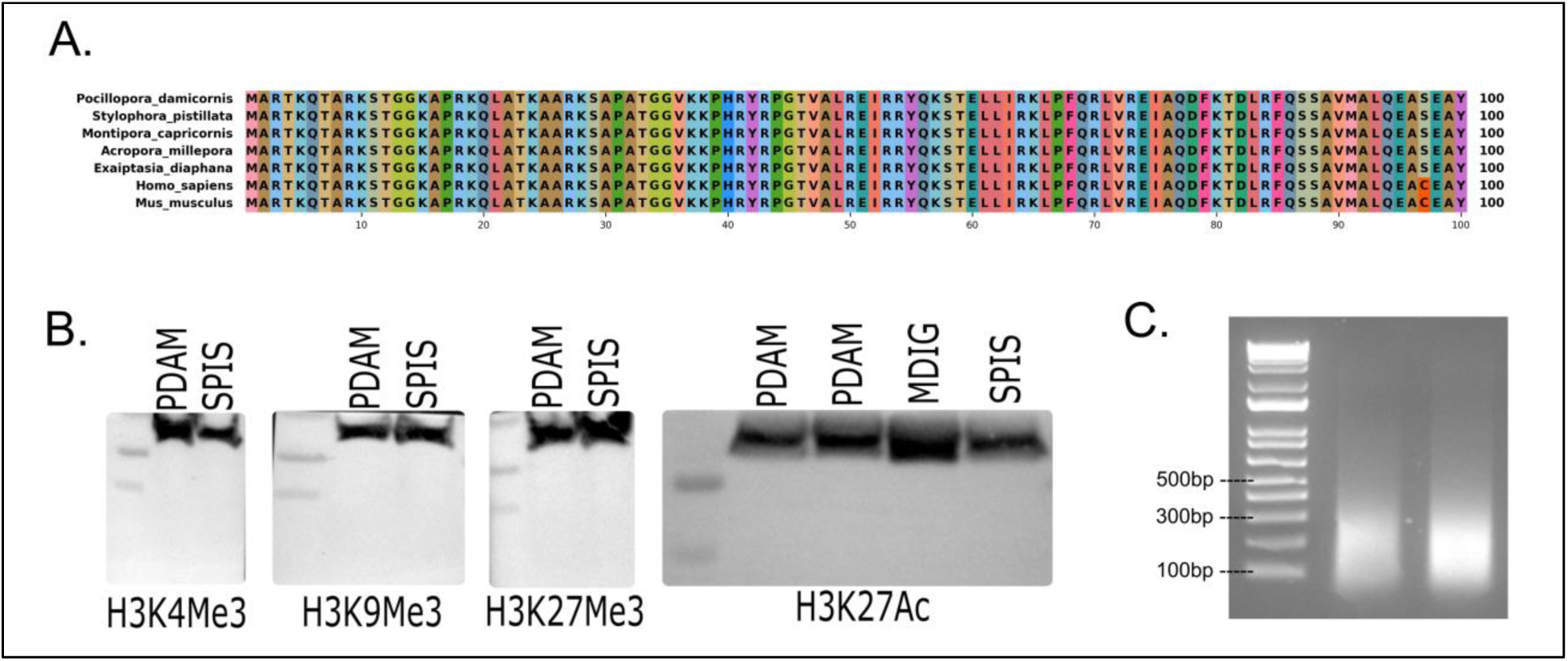
Protein alignment, antibody validation, and chromatin sonication. A) Protein alignment of histone H3 protein across several reef-building corals, *Aiptasia* anemone, mouse, and human genomes. Accessions: *P. damicornis* **LOC113680858**, *Stylophora pistillata* **LOC111340752,** *Montipora capricornis* **LOC138038771,** *Acropora millepora* **LOC114952568,** *Exaiptasia diaphana* **LOC110234914,** *Mus musculus* **NM_145073.2,** *Homo sapiens* **NM_003536.3**. Protein homologs demonstrate near-perfect residue conservation. B) Western blots of four canonical H3 histone modifications for protein extracts from three different reef-building coral species: *Pocillopora damicornis* (PDAM), *Stylophora pistillata* (SPIS), and *Montipora digitata* (MDIG). Ladder markings below protein bands are 15kDa and 10kDa respectively. C) Electrophoresis gel image of chromatin shearing in the input controls (whole cell extracts) for the two pooled chromatin replicates used in this study.

**Figure S2.**
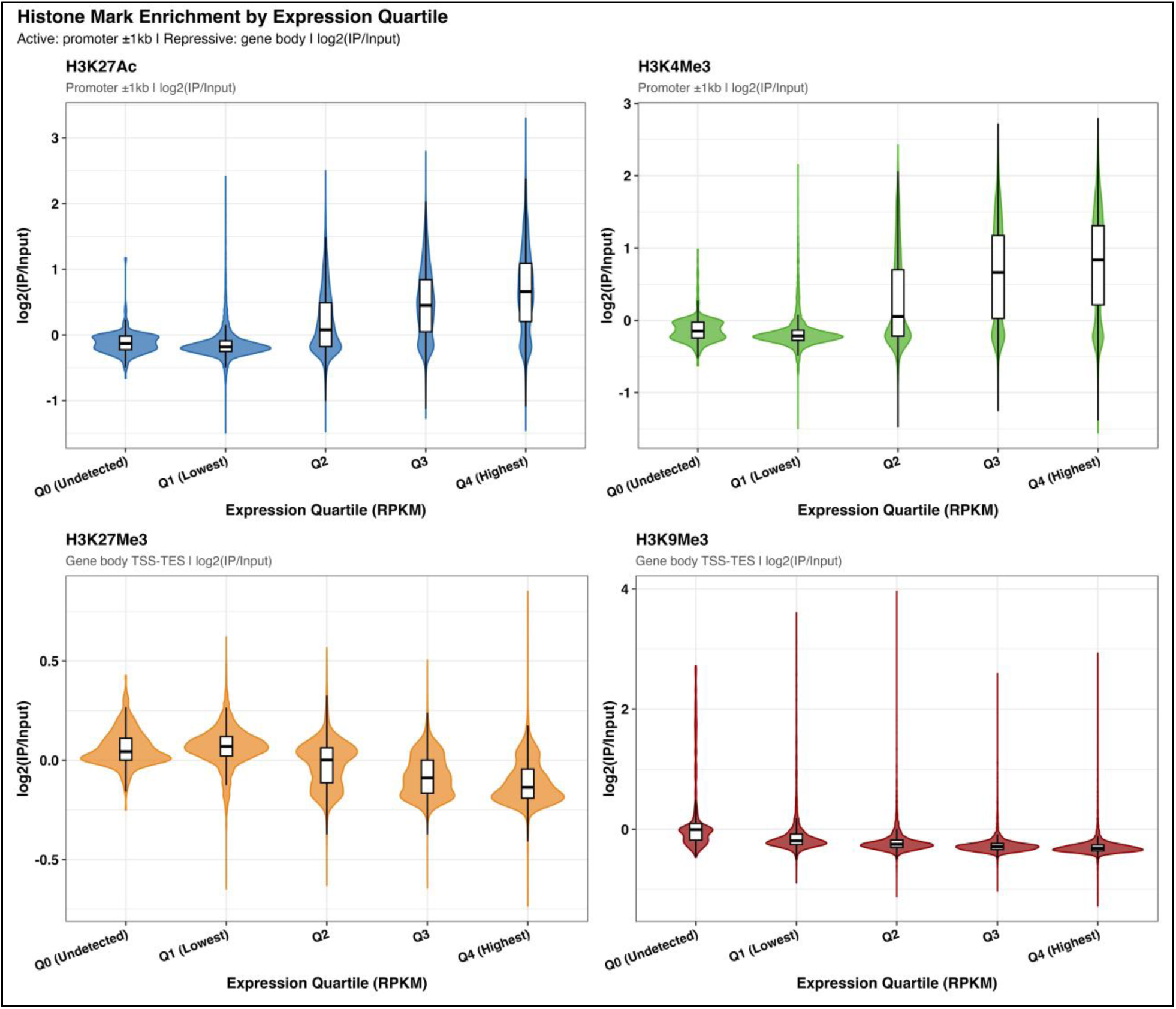
Violin plots of histone modification scores across expression quartiles. Mean log2(IP/input) ChIP signal in promoters (for H3K4me3 and H3K27ac) and in gene bodies (for H3K27me3 and H3K9me3) across expression groups as partitioned by RNA-seq RPKM counts, to accompany Figure 2A.

**Figure S3.**
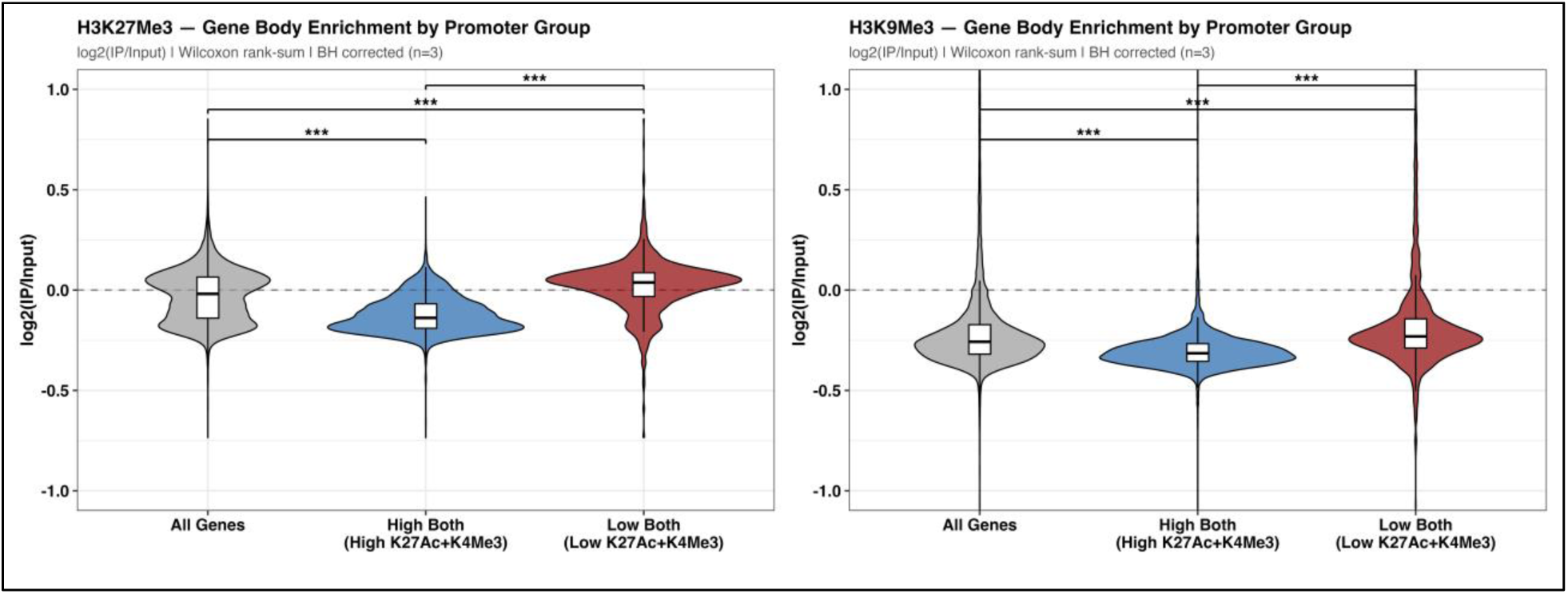
Repressive mark gene body score distributions in promoter classes. Mean log2(IP/input) H3K27me3 (*left)* and H3K9me3 (*right)* ChIP signal in gene bodies across all RefSeq genes, High, and Low promoter group genes. Significance indicated by Wilcoxon Rank-sum tests, BH-corrected for n=3 tests (p<0.001, ***).

**Table S1.** Statistical testing of log2(IP/input) histone mark scores across gene expression quartiles. Kruskal Wallis Dunn pairwise statistics (with BH correction) for violin/gene expression plots in Figure 2A and Figure S2.

| Mark | Group 1 | Group 2 | Z | p_value | p_adj | Sig Stars |
| --- | --- | --- | --- | --- | --- | --- |
| H3K27ac | Q0 (Undetected) | Q1 (Lowest) | 1.88 | 6.01E-02 | 6.01E-02 | ns |
| H3K27ac | Q1 (Lowest) | Q2 | -36.334 | 4.72E-289 | 1.18E-288 | *** |
| H3K27ac | Q2 | Q3 | -25.385 | 3.74E-142 | 7.47E-142 | *** |
| H3K27ac | Q3 | Q4 (Highest) | -11.756 | 6.60E-32 | 8.25E-32 | *** |
| H3K27me3 | Q0 (Undetected) | Q1 (Lowest) | -1.45 | 1.47E-01 | 1.47E-01 | ns |
| H3K27me3 | Q1 (Lowest) | Q2 | 34.982 | 4.21E-268 | 1.05E-267 | *** |
| H3K27me3 | Q2 | Q3 | 24.896 | 8.26E-137 | 1.65E-136 | *** |
| H3K27me3 | Q3 | Q4 (Highest) | 13.325 | 1.65E-40 | 2.06E-40 | *** |
| H3K4me3 | Q0 (Undetected) | Q1 (Lowest) | 2.88 | 3.98E-03 | 3.98E-03 | ** |
| H3K4me3 | Q1 (Lowest) | Q2 | -36.108 | 1.71E-285 | 5.69E-285 | *** |
| H3K4me3 | Q2 | Q3 | -26.313 | 1.35E-152 | 2.69E-152 | *** |
| H3K4me3 | Q3 | Q4 (Highest) | -6.949 | 3.67E-12 | 4.08E-12 | *** |
| H3K9me3 | Q0 (Undetected) | Q1 (Lowest) | 7.677 | 1.62E-14 | 1.62E-14 | *** |
| H3K9me3 | Q1 (Lowest) | Q2 | 24.983 | 9.45E-138 | 1.89E-137 | *** |
| H3K9me3 | Q2 | Q3 | 22.045 | 1.08E-107 | 1.54E-107 | *** |
| H3K9me3 | Q3 | Q4 (Highest) | 13.383 | 7.57E-41 | 8.42E-41 | *** |

**Table S2.** Gene expression data for neurotransmitter-like genes. RPKM counts of identified neurotransmitter-like genes that exhibit H3K27me3 repression in Figure 4.

| NCBI ID | Product Description | Average RPKM Counts |
| --- | --- | --- |
| 113682779 | octopamine receptor-like | 0.947 |
| 113664240 | histamine H2 receptor-like | 0.494 |
| 113664213 | beta-1 adrenergic receptor-like | 1.512 |
| 113664214 | beta-1 adrenergic receptor-like | 0.293 |
| 113666351 | D(2) dopamine receptor-like | 0.343 |
| 113666348 | beta-1 adrenergic receptor-like | 0.826 |
| 113666353 | beta-1 adrenergic receptor-like | 0.240 |
| 113666648 | alpha-1B adrenergic receptor-like | 0.477 |
| 113670560 | D(1)-like dopamine receptor | 0.088 |
| 113672888 | beta-1 adrenergic receptor-like | 1.506 |
| 113672863 | 5-hydroxytryptamine receptor 1A-like | 0.417 |
| 113673516 | probable G-protein coupled receptor No9 | 0.042 |
| 113677540 | D(2) dopamine receptor A-like | 0.143 |
| 113680416 | histamine H2 receptor-like | 1.222 |
| 113683128 | histamine H2 receptor-like | 0.225 |
| 113683397 | histamine H2 receptor-like | 0.544 |
| 113683601 | probable G-protein coupled receptor No9 | 7.372 |
| 113684880 | octopamine receptor beta-2R-like | 1.471 |

**Table S3.**
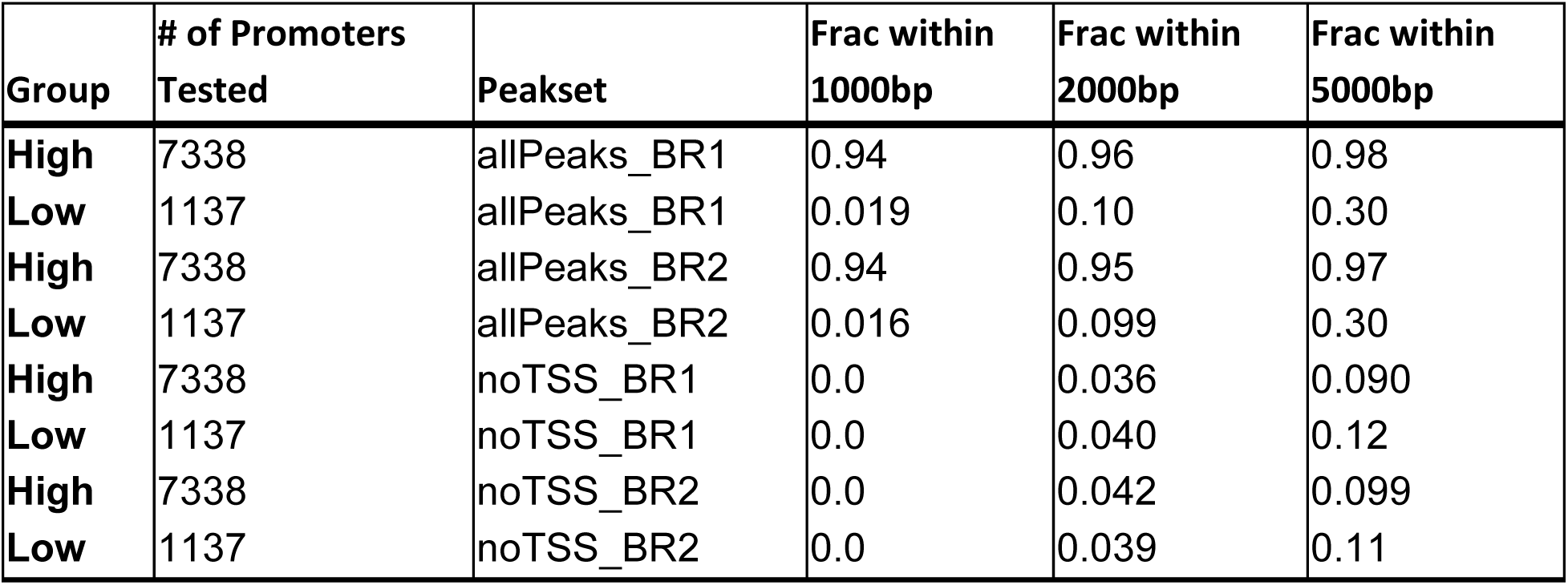
TSS misannotation sensitivity analysis results. Genomic distances between the Refseq annotated TSS in promoters and the nearest H3K4me3 peak are reported. Peaksets containing either 1) all *macs2* identified peaks, “allPeaks” or 2) only peaks not occurring in RefSeq annotated TSSs, “noTSS” (highest potential as misannotated TSSs). Promoter sets are the same as those used for motif discovery in Figure 3, where a small number of promoter sequences that occurred at the edges of scaffolds (where a full ±1kb from TSS was not possible) were excluded from the analysis (High: n=80, 1.1%; Low: n=136, 10.7%).

| Group | # of Promoters Tested | Peakset | Frac within 1000bp | Frac within 2000bp | Frac within 5000bp |
| --- | --- | --- | --- | --- | --- |
| High | 7338 | allPeaks_BR1 | 0.94 | 0.96 | 0.98 |
| Low | 1137 | allPeaks_BR1 | 0.019 | 0.10 | 0.30 |
| High | 7338 | allPeaks_BR2 | 0.94 | 0.95 | 0.97 |
| Low | 1137 | allPeaks_BR2 | 0.016 | 0.099 | 0.30 |
| High | 7338 | noTSS_BR1 | 0.0 | 0.036 | 0.090 |
| Low | 1137 | noTSS_BR1 | 0.0 | 0.040 | 0.12 |
| High | 7338 | noTSS_BR2 | 0.0 | 0.042 | 0.099 |
| Low | 1137 | noTSS_BR2 | 0.0 | 0.039 | 0.11 |

**Table S4.** Raw sequencing and mapping statistics for ChIP-seq libraries. BR# indicates the replicate number, and A/B indicates repeat immunoprecipitation of the same replicate. PDAM=*Pocillopora damicornis*, *C. gor*=*Cladocopium goreaui*, *D. tren*=*Durusdinium trenchii*.

| Library | # of Read Pairs | % Mapped to PDAM MapQ>30 | % Mapped to *C. gor* MapQ>30 | % Mapped to *D. tren* MapQ>30 |
| --- | --- | --- | --- | --- |
| H3K27ac BR1 | 51376466 | 74.9 | 2.83 | 0.055 |
| H3K27ac BR1B | 58775904 | 79.6 | 2.51 | 0.068 |
| H3K27me3 BR1 | 25043173 | 71.7 | 3.79 | 0.084 |
| H3K4me3 BR1 | 23545239 | 82.5 | 1.85 | 0.067 |
| H3K9me3 BR1 | 27140562 | 65.3 | 1.90 | 0.144 |
| Input Control BR1A | 32828305 | 78.7 | 1.96 | 0.093 |
| Input Control BR1B | 43086490 | 79.7 | 1.55 | 0.087 |
| H3K27ac BR2 | 31048564 | 78.8 | 3.07 | 0.068 |
| H3K27me3 BR2 | 32700915 | 74.5 | 2.98 | 0.082 |
| H3K4me3 BR2 | 30572853 | 80.6 | 2.06 | 0.067 |
| H3K9me3 BR2 | 41934328 | 67.8 | 2.00 | 0.123 |
| Input Control BR2A | 25976894 | 77.4 | 2.07 | 0.098 |
| Input Control BR2B | 46364998 | 79.7 | 1.73 | 0.092 |

**Table S5.**
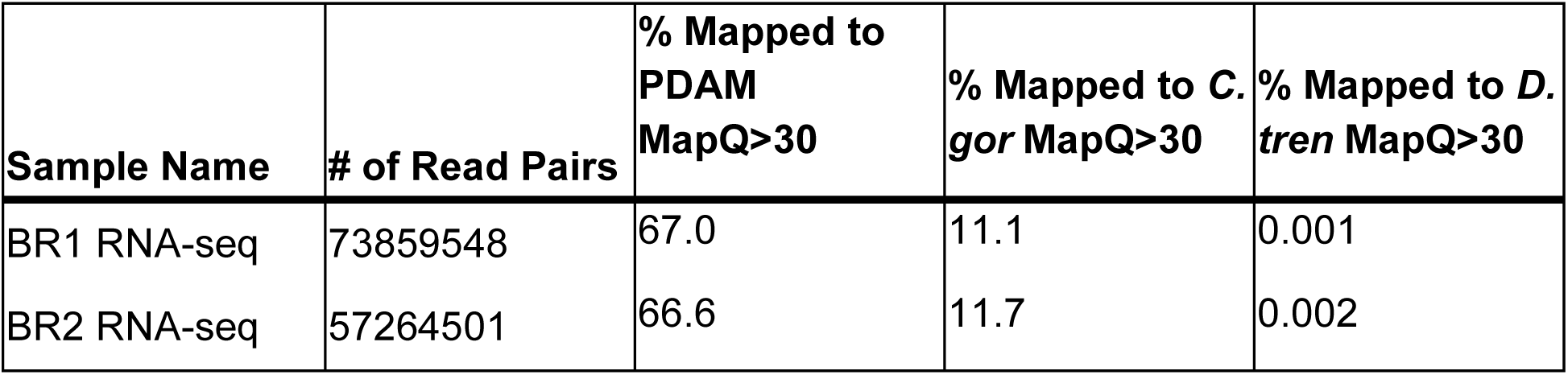
Raw sequencing and mapping statistics for RNA-seq libraries for the two tissue samples used in this study. . PDAM=*Pocillopora damicornis*, *C. gor*=*Cladocopium goreaui*, *D. tren*=*Durusdinium trenchii*.

| **Sample Name** | **# of Read Pairs** | **% Mapped to PDAM MapQ>30** | **% Mapped to *C. gor* MapQ>30** | **% Mapped to *D. tren* MapQ>30** |
| --- | --- | --- | --- | --- |
| BR1 RNA-seq | 73859548 | 67.0 | 11.1 | 0.001 |
| BR2 RNA-seq | 57264501 | 66.6 | 11.7 | 0.002 |

