## Supplementary Data 1 - STREME High Promoters (scrambled background) for "Genome-Wide Mapping of Major Histone Modifications Reveals Distinct Epigenetic Regulatory States in a Reef-Building Coral": Supplementary Data 1 - STREME High Promoters (scrambled background).html

STREME Results


Help poup.

[close ]

Motifs discovered by STREME in MEME motif format.

[
close ]

STREME results in XML format.

[
close ]

[
close ]

Read more about the MEME Suite's use of the IUPAC alphabets.

[close ]

[close ]

[close ]

[close ]

[close ]

[
close ]

[
close ]

The number of positive sequences matching the motif (percentage).

[close ]

The number of training set positive sequences matching the motif / the number of training set positive sequences.

Note these counts are made after erasing sites that match previously
found motifs.

[close ]

The number of training set positive sequences matching the motif.

Note these counts are made after erasing sites that match previously
found motifs.

[close ]

The number of training set negative sequences matching the motif / the number of training set negative sequences.

Note these counts are made after erasing sites that match previously
found motifs.

[close ]

The number test set positive sequences matching the motif / the number of test set positive sequences.

Note these counts are made after erasing sites that match previously
found motifs.

[close ]

The number of test set positive sequences matching the motif.

Note these counts are made after erasing sites that match previously
found motifs.

[close ]

The number of test set negative sequences matching the motif / the number of test set negative sequences.

Note these counts are made after erasing sites that match previously
found motifs.

[close ]

The mean distance from the center of the best match to the sequence center,
averaged over all training set sequences with a match.

[close ]

The mean distance from the center of the best match to the sequence center,
averaged over all test set sequences with a match.

[close ]

For determining if a motif is statistically significant, you should use the
value in the E-value column. If there is no E-value column, that
means that either the positive or negative hold-out set would have been too small
(fewer than 5 sequences).
For very small sequence sets, it is not practical for STREME to compute an accurate *E*-value.
In that case, you can determine if your motif is significant by running STREME twenty or more
times on shuffled versions of your positive dataset,
and seeing if the Score is always **larger** than the Score using the original sequences.
You can make shuffled sequence datasets using the MEME Suite command-line utility
fasta-shuffle-letters) if you
have installed the MEME Suite on your own computer.

The statistical test used in computing the Score is either the Fisher Exact Test,
the Binomial Test, or the Cumulative Bates distribution. (See Inputs and Settings
for the particular test being used.) The Fisher Exact Test and the Binomial Test both
estimate the enrichment of the motif in the positive sequences compared to the
the negative sequences.
(The Binomial Test is used when the positive and negative sequences have different average lengths.)
The Cumulative Bates distribution measures the tendency
of motif to be near the center of the input sequences.

[close ]

The statistical test used in computing the *p*-value is either the Fisher Exact Test,
the Binomial Test, or the Cumulative Bates distribution. (See Inputs and Settings
at the bottom of this document for the particular test being used.)
The Fisher Exact Test and the Binomial Test both
measure the enrichment of the motif in the positive test sequences compared to the
the negative test sequences.
(The Binomial Test is used when the positive and negative sequences have different average lengths.)
The Cumulative Bates distribution measures the tendency
of motif to be near the center of the sequences.

[close ]

[close ]

The score threshold for determining if a potential site is a match
to the motif. The same threshold is applied when determining matches in
the training and test sequences. The threshold is in bits.

The match score of a position in a sequence is determined by converting
the motif to a base-2 log-odds matrix using the formula log2(prob[a][i]/background[a]).
Here, prob[a][i] is the probability of the letter 'a' at position 'i' of the motif,
and background[a] is the probability of the letter 'a' according to the background.

[close ]

The names of the files containing the positive (primary)
and negative (control) sequences input to STREME.

If you did not provide a file containing the negative (e.g., control)
sequences, STREME created them using N-order shuffling.
0-order shuffling preserves 1-mer frequencies (i.e., the letter frequencies),
1-order shuffling preserves 2-mer frequencies, etc.

[close ]

The name of the alphabet of the sequences.

[close ]

The number of sequences.

[close ]

The total length of the sequences.

[close ]

The name of the alphabet symbol.

[close ]

The frequency of the alphabet symbol in the negative sequences.

[close ]

The frequency of the alphabet symbol as defined by the background model.

[
close ]

###### Details

| Train Positives | Train Positives | Train Negatives | Train DTC | Score | Test Positives | Test Positives | Test Negatives | Test DTC | P-value | Match Threshold |
| --- | --- | --- | --- | --- | --- | --- | --- | --- | --- | --- |
| /  () | /  () |  |  |  | /  () | /  () |  |  |  |  |

### STREME

#### Sensitive, Thorough, Rapid, Enriched Motif Elicitation

For further information on how to interpret these results please access
https://meme-suite.org/meme/doc/streme.html.   
To get a copy of the MEME software please access
https://meme-suite.org.

Discovered Motifs
  |  
Inputs & Settings
  |  
Program Information
  |  
Motifs in MEME Text Format 
  |  
Matching Sequences >
  |  
Results in XML Format


### Javascript is required to view these results!

### Your browser does not support canvas!


#### Discovered Motifs

Next Top

No motifs were discovered!

#### Inputs & Settings

Previous Next Top

###### Sequences

| Role | Source | Alphabet | Sequence Count | Total Size |
| --- | --- | --- | --- | --- |
| Positive (primary) Sequences |  |  |  |  |
| Negative (control) Sequences |  |  |  |  |

###### Background Model


###### Other Settings

|  |  |
| --- | --- |
| Strand Handling | This alphabet only has one strand. Only the given strand is processed. Both the given and reverse complement strands are processed. |
| Objective Function |  |
| Statistical Test |  |
| Motif Selection Criterion |  |
| Minimum Motif Width |  |
| Maximum Motif Width |  |
| Sequence Shuffling |  |
| Test Set |  |
| Word Evaluation |  |
| Seed Refinement |  |
| Refinement Iterations |  |
| Minimum Score |  |
| Refinement Match Subsets |  |
| Minimum Palindrome Ratio |  |
| Maximum Palindrome Edit Distance |  |
| Print Candidate Motifs? |  |
| Random Number Seed |  |
| Total Length |  |
| Maximum Motif |  |
| Maximum Motifs to Find |  |
| Maximum Run Time |  |

Previous Top

###### STREME version

(Release date: )

###### Command line
